# Enhancement of hidden Markov model analyses for improved inference of archaic introgression in modern humans

**DOI:** 10.1101/2025.04.22.649993

**Authors:** Moisès Coll Macià, Laurits Skov, Zenia Elise Damgaard Bæk, Asger Hobolth

## Abstract

Insights into the admixture history between modern and archaic humans require accurately inferred introgressed fragments within modern genomes. Here, we introduce two enhancements to hidden Markov models (HMMs) implemented in hmmix. First, we develop a method for sampling hidden state sequences conditional on observed genomic data, enabling robust estimation of admixture summary statistics—such as admixture proportion and fragment length distributions. This represents an improvement compared to relying solely on point estimates as provided by classical decoding methods. Additionally, we integrate the Finite Markov Chain Imbedding (FMCI) framework, allowing exact analytical calculation of these admixture statistics, tailored to large scale human genomes. Second, we implement a novel hybrid decoding method which combines the strengths of Viterbi and Posterior decoding methods, substantially improving the reliability of archaic fragments identified. We validate these improvements on data from the 1000 Genomes Project and demonstrate that our sampling method yield more accurate admixture estimates from single individuals compared to existing approaches requiring extensive population-level datasets. Moreover we show how hybrid decoding can be instrumental in resolving the inference of local archaic haplotype structure in modern human genomes. These methodological advancements broadly enhance HMM-based analyses and will provide deeper insight into the complex history of genetic interactions between archaic and modern human populations.

## Introduction

Decoding archaic content in modern humans provides insights into past interactions between our ancestors and the extinct Neanderthals and Denisovans (Iasi et al. 2024; Skov et al. 2020; Bergström et al. 2020; Sankararaman et al. 2016). A key tool in this effort is hmmix (Skov et al. 2018), which identifies archaic fragments in human genomes based on Hidden Markov Models (HMMs). HMMs are well-suited for this task due to the sequential nature of genomic data and because archaic ancestry cannot be directly measured, but must be inferred from indirect genetic signatures.

In general, Posterior decoding and Viterbi decoding are the main methods to decode the hidden state sequences using HMMs (Zucchini, MacDonald and Langrock 2017). For the case of hmmix, Posterior decoding is the default option as its results are more accurate due to its definition of optimizing local probabilities by maximizing accuracy in each position of the sequence. In contrast, Viterbi decoding is the hidden path with the highest probability among all possible hidden paths conditional on the observed sequence. This approach optimizes a global property and the resulting hidden path has substantially fewer hidden state transitions than Posterior decoding. Although this behaviour reduces its risk for detecting false archaic fragments (false positives), it makes the model miss short yet genuine archaic fragments.

Despite their respective strengths, each decoding method represents extreme optimization strategies, either local or global, without possible intermediate solutions. Another limitation of the existing decoding methods is that both produce deterministic outputs, yielding a single fixed set of archaic fragments per genome without an explicit quantification of uncertainty.This lack of uncertainty estimation can bias downstream analyses based on summary statistics, such as estimates of admixture proportions, potentially leading to inaccurate interpretations of archaic ancestry in the genome analysed.

To address these issues, we describe two key enhancements for general HMM frameworks in a separate manuscript (Bæk et al. 2025), and here tailored to hmmix. First, we use the fact that the hidden state sequence, conditional on the observed sequence, is an inhomogeneous Markov chain. The inhomogeneous transition probabilities along the sequence can be found from the backward table of the HMM. The inhomogeneous transition probabilities enable estimating the full distributions for the summary statistics of interest with a sampling based as well as an analytical based approach (Aston and Martin 2007). From this, we obtain a more robust and accurate estimation of key parameters in the study of archaic admixture, such as the distributions of archaic fragment length. Second, we implement the hybrid decoding method (Lember and Koloydenko 2014; Kuljus and Lember 2023). This method estimates hidden state sequences that are intermediate solutions between Posterior and Viterbi outputs by combining the strengths of both global and local decoding optimizations, respectively. Finally, we apply these methodological enhancements to three real phased genomes. We demonstrate that the new methodologies allow us to obtain more refined insights about the admixture history between modern and archaic humans, showcasing the improvements that can be obtained when applying these approaches to HMM analyses.

## Results

### hmmix

Non-African human genomes carry ∼2% to 5% Neanderthal and Denisovan DNA, a legacy from the encounters between archaic hominins and early modern humans during their expansion out of Africa (Bergström et al. 2020). As a result, non-Africans have accumulated private genetic variants in two distinct ways when compared to their African counterparts: through new mutations that arose after the divergence from African groups, and through archaic admixture. Notably, archaic fragments in modern human genomes exhibit a much higher density of private variants, proportional to the longer evolutionary timespan to accumulate those (Figure 1).

**Figure 1.**
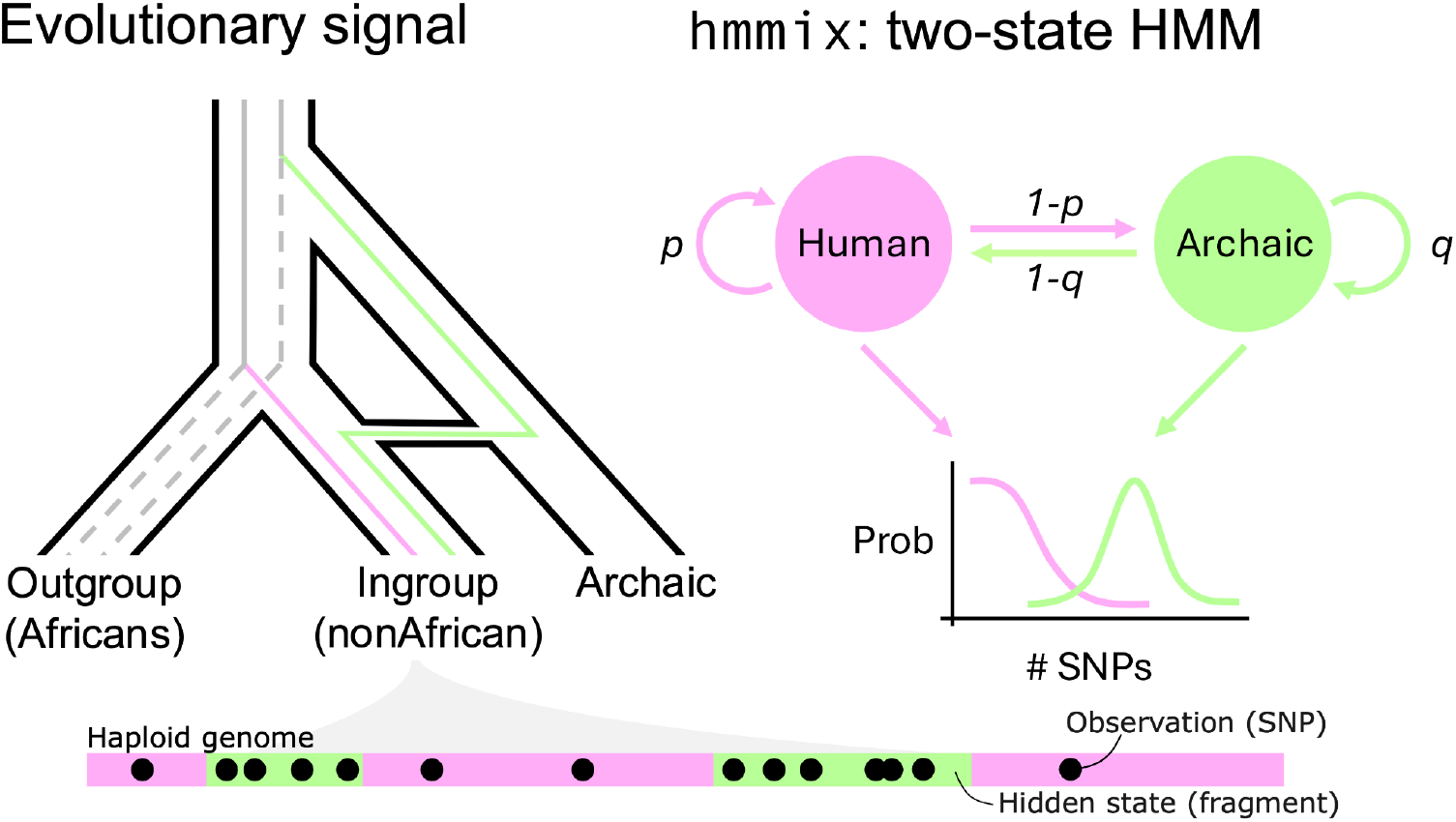
Graphical representation of the evolutionary signal (left) that hmmix uses as input and the model itself (right). The tree shows the phylogenetic relationship between the outgroup, ingroup and the archaic group. Only the ingroup receives gene-flow from the archaic group. The lines within the tree represent archaic (green) and modern human (pink) lineages within the ingroup genome. Colorful parts of the lineages show the time while each lineage accumulates private genetic variation compared to the outgroup. A representation of a haploid genome from an ingroup individual is represented as a strip with human and archaic fragments. Black dots represent private variation accumulated in those regions, with archaic fragments being more dense than human fragments. The graph on the right represents hmmix as the two-state HMM with their transition probabilities between archaic and human states and their corresponding emission probabilities with different Poisson rates depending on SNP density.

The hmmix software (Skov et al. 2018) leverages this contrast of private SNP density to identify genomic regions of archaic origin in test genomes (Figure 1). It employs a two-state HMM in which the hidden states represent archaic and human genetic origin. Observed SNP densities in 1000 bp (1 kb) windows are modeled as Poisson-distributed, with different rates based on the underlying state. To achieve this difference of SNP density in genomic data, common variants segregating in a set of African genomes (outgroup) are excluded from the test genome (ingroup). Compared to similar methods, this design enables hmmix to infer archaic fragments without requiring a reference archaic genome, thereby reducing potential bias towards any specific sequenced Neanderthal or Denisovan genome.

To evaluate the advances implemented in hmmix, we simulate archaic fragments under two distinct parameter sets with hmmix (Methods). The first set, referred to as realistic parameters, mimics the Neanderthal admixture scenario and is used for the statistical evaluation of the different methods presented (Table 1). The second parameter set, named as visualization parameters, produces a much higher overall archaic content and is designed mainly for visualization purposes and facilitating interpretation (Table 2).

**Table 1.** Realistic parameters and their Baum-Welch estimated parameters with their 95% CI (Methods). The estimated parameters are rounded to the fourth digit.

| Realistic parameters |  |  |  |  |
| --- | --- | --- | --- | --- |
| | Emission probabilities ($\lambda$) | Starting probabilities ($\pi$) | Transition probabilities ($\Gamma$) | |
|  |  |  | Human | Archaic |
| Human | 0.03 | 0.98 | 0.9995918 | 0.0004082 |
| Archaic | 0.3 | 0.02 | 0.02 | 0.98 |

| Baum-Welch estimated realistic parameters |  |  |  |  |
| --- | --- | --- | --- | --- |
| | Emission probabilities ($\lambda$) | Starting probabilities ($\pi$) | Transition probabilities ($\Gamma$) | |
|  |  |  | Human | Archaic |
| Human | 0.0300<br>(0.0299 - 0.0302) | 0.9808<br>(0.9796 - 0.9820) | 0.9996<br>(0.9996 - 0.9996) | 0.0004<br>(0.0004 - 0.0004) |
| Archaic | 0.3039<br>(0.3000 - 0.3090) | 0.0192<br>(0.0180 - 0.0204) | 0.0204<br>(0.0191 - 0.0215) | 0.9796<br>(0.9785 - 0.9809) |

**Table 2.** Visualization parameters and their Baum-Welch estimated parameters.

| Visualization parameters |  |  |  |  |
| --- | --- | --- | --- | --- |
| | Emission probabilities ($\lambda$) | Starting probabilities ($\pi$) | Transition probabilities ($\Gamma$) | |
|  |  |  | Human | Archaic |
| Human | 0.2 | 0.9 | 0.99 | 0.01 |
| Archaic | 0.5 | 0.1 | 0.02 | 0.98 |

| Baum-Welch estimated visualization parameters |  |  |  |  |
| --- | --- | --- | --- | --- |
| | Emission probabilities ($\lambda$) | Starting probabilities ($\pi$) | Transition probabilities ($\Gamma$) | |
|  |  |  | Human | Archaic |
| Human | 0.200 | 0.674 | 0.990 | 0.010 |
| Archaic | 0.500 | 0.327 | 0.020 | 0.980 |

### Summary distributions from the conditional hidden state sequence

The hidden state sequence in a HMM, conditioned on the observed data, constitutes an inhomogeneous Markov chain. The transition probabilities between the hidden states become inhomogeneous because they depend on the observed data (Methods, Bæk et al. 2025). In other words, the probability of staying in the same state or changing to another state varies throughout the sequence depending on the observations. (Aston and Martin 2007) discussed this property, highlighting its potential for sampling from the conditional distribution of state sequences. This approach thus can generate a collection of simulated hidden state sequences, reflecting the variance in the conditional posterior probabilities.

Building on this idea, we implement in hmmix a procedure for sampling from the inhomogeneous Markov chain of hidden state sequences conditional on the observed data (Methods). In Figure 2a we show a simulated observed sequence from the visualization parameters (Methods), and in Figure 2b we show the corresponding inhomogeneous transition probabilities of staying in each state conditional on the observed sequence (Methods). The probabilities of staying in the archaic state *b*_*t*_ increase with the number of observed variants per window. Conversely, the probabilities of staying in the human state *a*_*t*_ decrease in the same regions. The reverse is true for low-density regions. This behavior aligns with the design of hmmix, where archaic regions are characterized by high variant density, while human regions correspond to lower densities (Figure 1).

**Figure 2.**
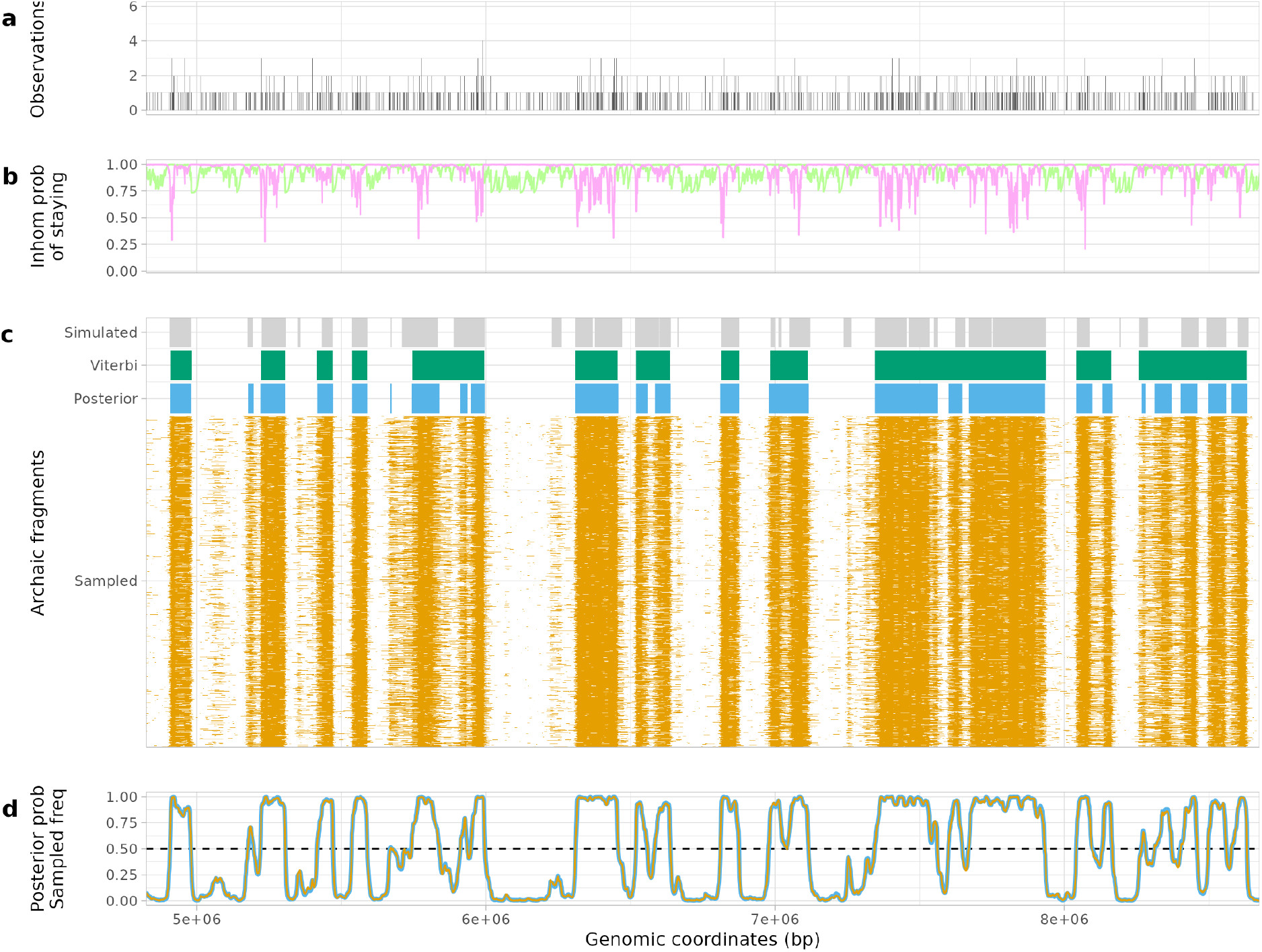
Simulated genetic region analysis from visualization parameters with results from different decoding methods. a) The observations as counts of SNPs in 1,000 bp windows as the input for hmmix. b) Inhomogeneous transition probabilities of staying in the same state (Archaic *b*_*t*_ in green and Human *a*_*t*_ in pink) conditional on the observed data. c) The archaic fragments from true (gray), Viterbi decoded (green), Posterior decoded (blue) and 1,000 samples from the conditional distribution of hidden states (orange). d) The posterior probability of being archaic from Posterior decoding (blue) and the frequency of archaic sequence in the 1,000 sampled paths (orange). Note that the two lines are superposed. The dashed black line denotes the posterior probability threshold (50%) to decide if a window is considered archaic or human for Posterior decoding.

Using the inhomogeneous transition probabilities, we sample 1,000 hidden state sequences for the visualization parameters’ simulated data with hmmix (Methods). These sequences are compared against the true simulated sequences, as well as those inferred by Viterbi and Posterior decoding implemented also in hmmix (Figure 2c).

Occasionally, sampling from the posterior is able to capture fragments missed by both Viterbi and Posterior decoding, proportional to their posterior probability. In the case of Posterior decoding, these fragments are missed because only fragments with posterior probabilities above 50% of being archaic are classified as such (Figure 2c,d). In fact, the frequency of a given position to be classified as archaic among all samples corresponds to its posterior probability as estimated by Posterior decoding (Pearson correlation test = 0.9997, p value < 2.2×10^-16^, Figure 2d).

The sampling approach allows us to compute the distribution of various summary statistics such as total amount of archaic sequence, archaic fragments length distribution, total number of archaic fragments and the longest archaic fragment. These statistics are relevant since in the study of archaic introgression, the total amount of archaic sequence in a sampled human genome is often used to estimate the admixture proportion from Neanderthals, for instance. Figure 3a shows three of the summary statistics obtained with the different decoding methods presented above and compares them to the true simulated value from realistic parameters. While only point estimates are obtained with the two classical decoding methods, with the sampling approach we are able to quantify the variance around the estimated summary of interest. In all three cases, the true value is captured by the distribution of summaries obtained with the 1,000 sampled paths. Conversely, both Viterbi and Posterior decoding are notably distant from the true values.

**Figure 3.**
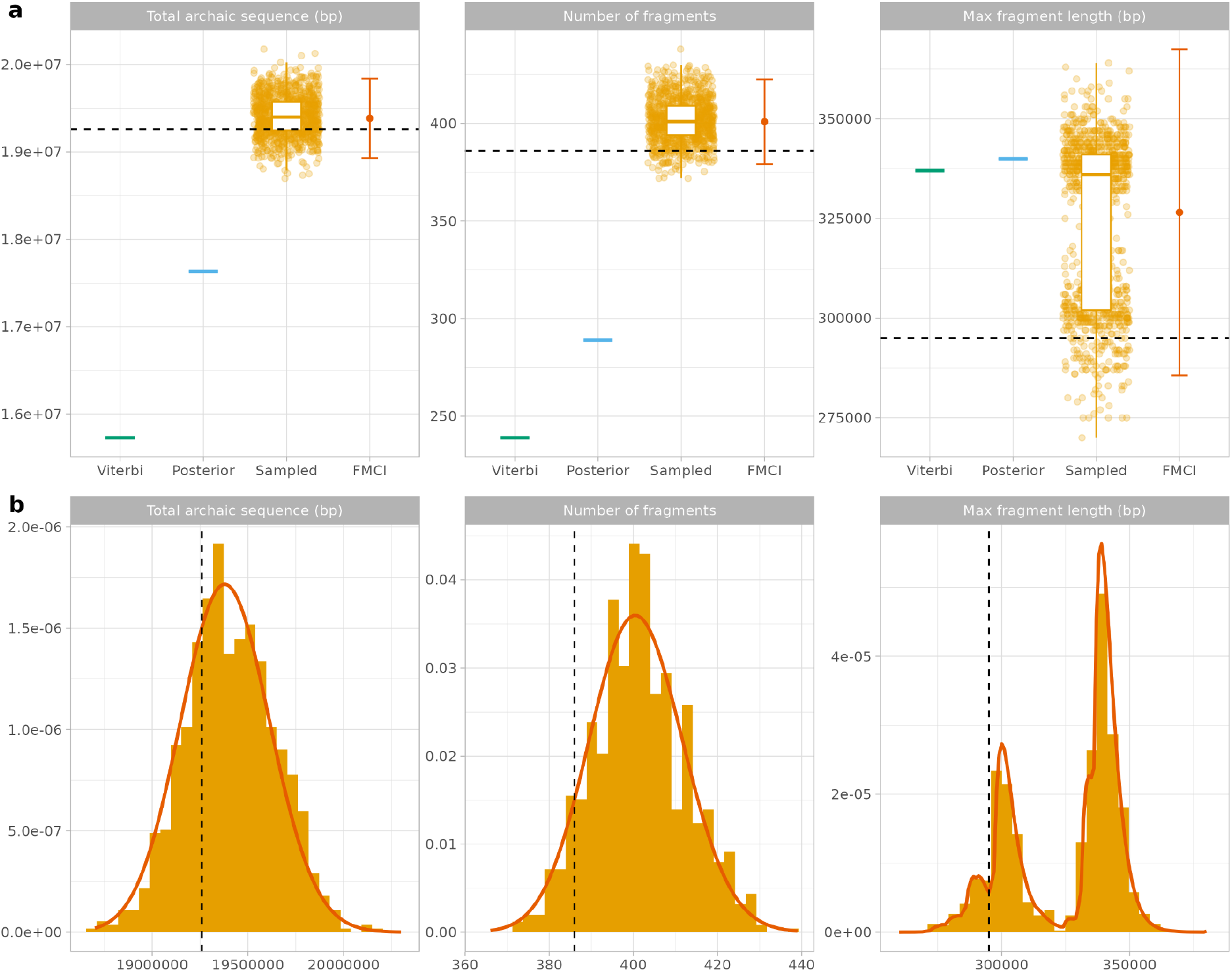
Summary statistics of archaic fragments based on realistic parameter simulations. a) Comparison between the true value form simulations (dash lines) with the summary obtained from each decoding method. The Viterbi (green) and Posterior decoding (blue) summary values are shown as horizontal lines. Individual values from each path sampled from the posterior (1,000 paths) are shown as orange dots and the distribution of all values are summarized with a box plot. The mean of the FMCI distribution is shown as a dark orange dot with the 1.96 standard deviation around the mean as error bars. b) Distribution of summary statistics from 1,000 paths sampled from the posterior (orange histogram) and the analytical FMCI distribution (dark orange line).

The accuracy estimating the summary distribution with this sampling based method is limited by the number of simulations performed. Instead, the analytical distribution of summary statistics can be computed using the Finite Markov Chain Imbedding (FMCI) (Aston and Martin 2007). This method consists of designing a first order Markov chain to compute the pattern of interest in which its transition probabilities are defined by the inhomogeneous transition probabilities introduced above (Methods, Supplementary Material 1, Bæk et al. 2025). The flexibility of this method to define any kind of Markov chain, enables us to build a framework to compute the distribution for the three summary statistics (Figure 3).

Depending on the specifics of each statistic, the FMCI varies in complexity, thus affecting the running time and memory efficiency of the algorithm (Supplementary Material 1). While computing the number of archaic fragments takes ∼20 minutes for the entire 1Gb-simulated sequence (Figure S1), it is infeasible to compute the distribution of the total amount of archaic sequence. Instead, we decided to divide the sequence into 100 10Mb chunks and compute the summary for each chunk in parallel. This way, we both reduce the maximum value to be computed (matrix size, *l*) and the sequence analyzed (number of iterations, *t*) ending up with a <2 min running time for each genomic chunk, which are able to run in parallel. To merge the results from each chunk together, we convolve the obtained distributions into a single one. To compute the maximum fragment length, we designed a different strategy. This is because the matrix computed scales quartically with the maximum length to be calculated (*l*^4^, Supplementary Material 1). To be able to run this algorithm, we compute sections of the distribution independently. We manage to make each job run in <4 hours. Finally, the resulting partial distributions are convolved together.

Both sampling from the posterior and FMCI methods can also recover specific features, such as the length distribution of archaic fragments (Figure 4, Bæk et al. 2025). In this case, both Posterior and Viterbi decodings also yield a distribution, but they are known to be biased since many small fragments are often missed. To compute the fragment length distribution with FMCI, we computed sections of the distribution independently due to the time complexity of the Markov chain (Supplementary Material 1). We also reused the same 1,000 samples from the posterior to calculate the distribution of interest. Figure 4 compares the true fragment length distribution with the ones estimated with the various methods. As expected, classical decoding methods in general struggle to recover short fragments, with Viterbi being exceptionally inferior at this. As a result, the mean fragment length is overestimated. In contrast, both the sampling method and FMCI more accurately reconstruct the true distribution with an unbiased mean.

**Figure 4.**
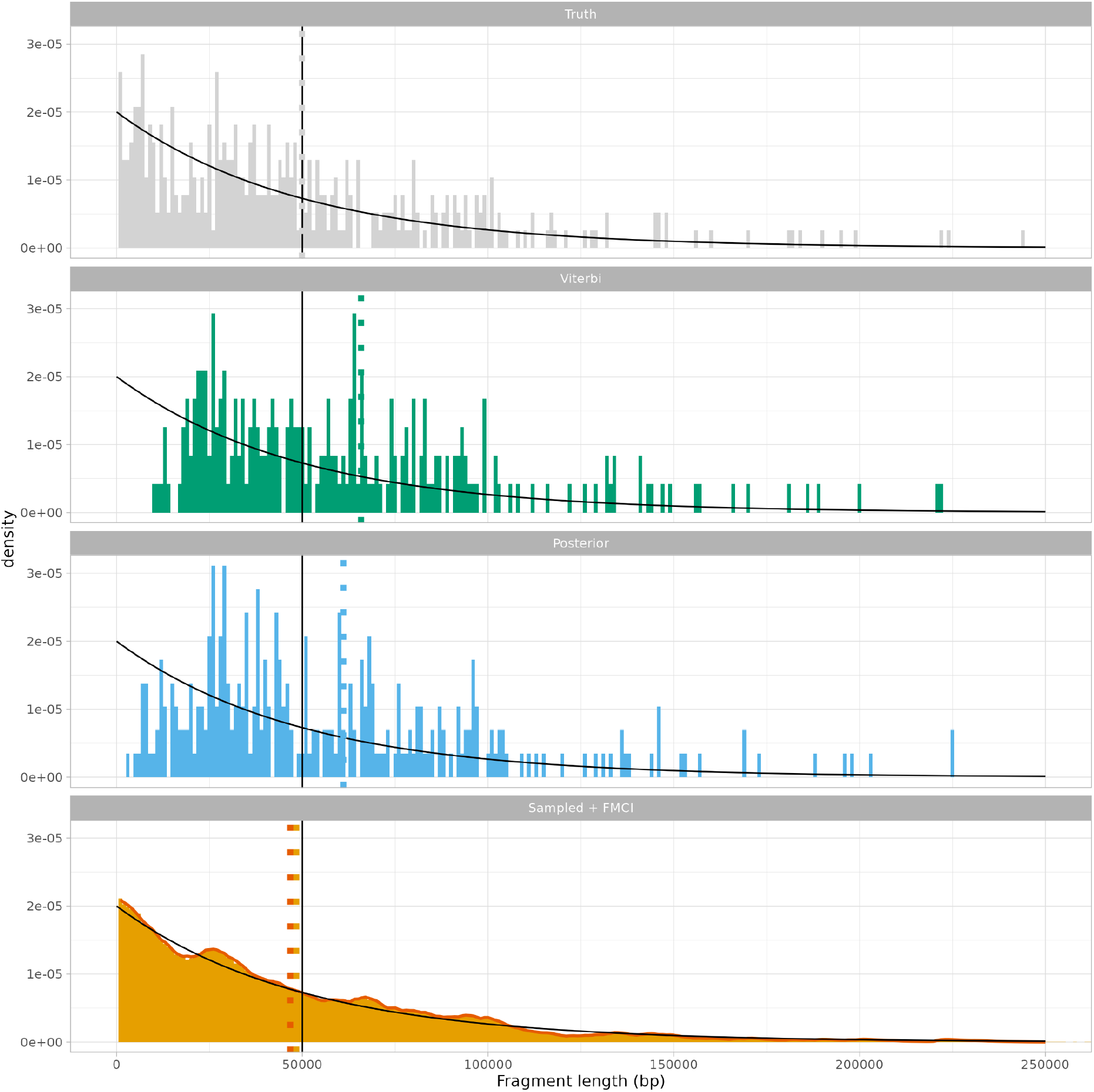
The length distribution of archaic fragments based on realistic parameter simulations for the true fragments (grey), the Viterbi (green) and Posterior decoded (blue) along with 1,000 paths sampled from the conditional distribution of hidden states (orange). The FMCI distribution is plotted on top of the sampled paths’ distribution (dark orange). Vertical dotted lines show the mean of each distribution with the same color code. The black decaying lines denote the theoretical length distribution of archaic fragments computed from a geometric distribution with parameter Γ_*AH*_ = 0. 02 and the vertical black lines its mean. The plot is limited to 250,000 bp on the x-axis.

Since the inhomogeneous Markov chain approaches are more influenced by the observations, we expect that they will be more resilient to violations of the HMM assumptions. Based on simulations, we conclude that this sampling approach can identify the correct summary statistics under realistic emission parameters of the human and archaic states of *λ*_*A*_ = 0. 3 and *λ*_*H*_ = 0. 03 (Supplementary Material 2). We also find that the sampling approach is robust to overdispersion of the observation data due to variation in coalescence times (Supplementary Material 3). Finally, we tested whether the sampling approach accurately recovers fragment length distributions when they deviate from the geometric assumption of HMMs, such as when recombination rates follow the human genetic map (Supplementary Material 4). We find that variation in the recombination rate across the genome leads to an underestimate of the fragment length unless emission parameters are very different (Supplementary Material 4).

In general, for demographic parameters that are compatible with a human-archaic introgression scenario, we find that the sampling approach produces robust estimates of the summary statistics and outperforms Viterbi and Posterior decoding even though the HMM assumptions are violated.

### Hybrid decoding

While our conditional probabilities approaches for estimating summary statistics offers an avenue for admixture parameter inferences, the precise decoding of archaic fragments remains essential for more localized analyses. For example, accurately determining whether a specific gene in modern human genomes has been inherited from an archaic lineage requires a reliable sequence decoding.

Previous work (Lember and Koloydenko 2014; Kuljus and Lember 2023) proposed a hybrid decoding method that combines the strengths of both Viterbi and Posterior decoding (Methods). We reinterpret the original method by calculating the weighted geometric mean between Viterbi and Posterior decoding, and parametrizing the weights of each term with *α* ∈ [0, 1] (Methods, Bæk et al. 2025). Specifically, when *α* = 1 the hybrid decoding outputs the Viterbi solution and when *α* = 0, the Posterior decoding solution. Hybrid decodings are obtained with intermediate *α* values (Methods).

We run hybrid decoding with hmmix’s implementation (Methods) on the visualization parameters’ simulated data with values of *α* ranging from 0 to 1 in increments of 0.1 units (Figure 5). In Figure 5, there are some examples in which Posterior decoding inferes multiple fragments that are also inferred by the hybrid method with small *α* values. Those sequences transition to more unified fragments in hybrid decodings with increasing *α* values. Eventually, the hybrid estimates converge to the Viterbi solution which often joins multiple true fragments into a single path, emphasizing the importance of choosing *α* adequately.

**Figure 5.**
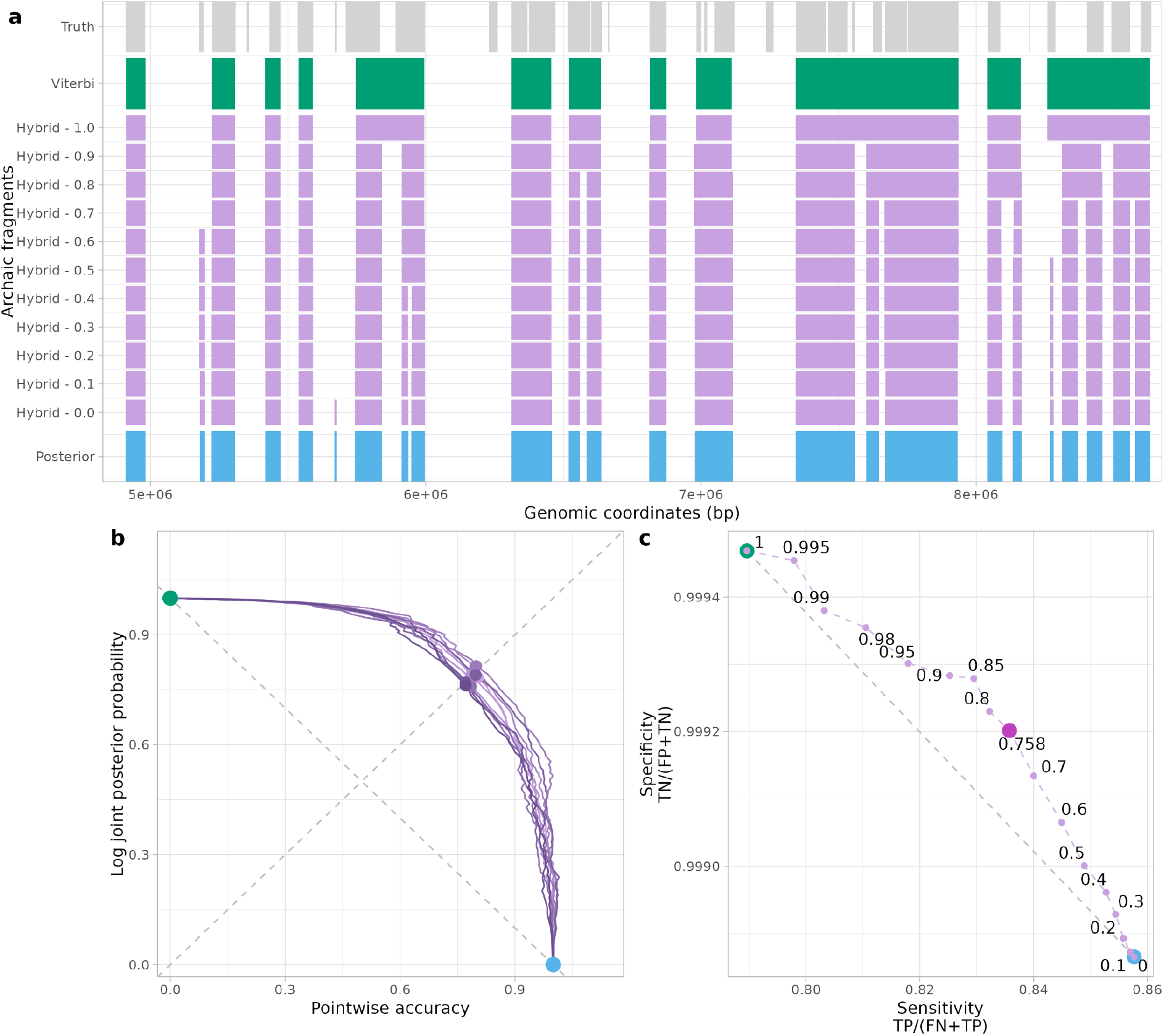
Hybrid decoding. a) Visualization of decoded paths using the hybrid method. The genomic region is the same as in Figure 2. The values of *α* are chosen to be between 0 (Posterior decoding) and 1 (Viterbi) in steps of 0.1. b) Artemis plot corresponding to the relationship between pointwise-accuracy and joined posterior probability of hybrid decoding paths in a grid search for *α* (min value = 0, max value = 1, step size = 0.01), the Viterbi and Posterior decoding paths (shown as dots in the extremes). 10 replicates are shown in this plot colored in different shades of purple. Optimal *α*’s are shown as dots for each replicate. c) Sensitivity (TN/(FP+TN)) and specificity (TP/(FN+TP)) values for the Viterbi, Posterior and hybrid paths for multiple *α* values. The optimal *α* value is shown as a bigger dot colored in magenta. Dashed line shows the range of values between Viterbi and Posterior decoding. TN : True Negatives; FP : False Positives; TP : True Positives; FN : False Negatives.

To tune *α*, we focus on the strengths of Viterbi and Posterior decoding: the log joint posterior probability and the pointwise accuracy, respectively. Since computing accuracy requires knowledge of the true sequence, tuning *α* relies on simulations. We simulate 10 hidden state sequences and the corresponding observables from the estimated parameters in the previous section (Table 1, Methods). We then perform a grid search of the parameter space of *α* in 0.01 increments (Figure 5b, Methods). We define an optimal *α* as the value whose hybrid solution is balanced in terms of the log joint posterior probability and the pointwise accuracy compared to the classical decoding methods, maximizing both simultaneously. In other words, the hybrid solution that lies closest to the 45º line between the maximum and minimum of both axes is considered optimal. This analysis – which we named Artemis due to the plot’s bow-shaped curvature and arrow-like 45º line – identifies an optimal *α* = 0.758 (95% CI 0.753 - 0.763) based on the 10 simulations of 10^9^ windows (Methods). Although the estimation of the optimal *α* has some variance among the 10 simulations, the resulting hybrid decodings using the extreme values within the 95% CI differ only marginally (Table S2), thus demonstrating the robustness of the parameter estimation with this method. Nonetheless, the hybrid method is flexible and *α* can be considered optimal depending on the user requirements (Bæk et al. 2025). We recommend running the Artemis analysis to optimize *α* in every case since the optimal point changes depending on the HMM parameters (Bæk et al. 2025). Artemis analysis is conveniently implemented in hmmix (Methods).

Next, we evaluate the performance of the hybrid method by assessing its specificity and sensitivity. As with the Artemis analysis, we estimated both metrics for hybrid decodings by performing a grid search over the parameter space—varying *α* in 0.1 increments, including additional values between 0.9 and 1 for finer resolution and the optimal *α* value 0.758. Figure 5c presents the full range of hybrid decoding results, which span between those of Posterior decoding and Viterbi. Notably, hybrid decodings lie above the grey dashed diagonal that connects the Posterior and Viterbi results; this diagonal represents an equal trade-off between specificity and sensitivity. Therefore, points above this line indicate better performance, maximizing both statistics simultaneously. The optimal hybrid path—previously identified in the Artemis analysis — corresponds to one of the points farthest from the diagonal, reflecting the best balance between sensitivity and specificity.

Overall, the hybrid implementation improves the hmmix archaic fragment inference by considering neighbouring signals in an intermediate scale between the local and global extreme optimization. This hybrid method reduces the relative proportion of false archaic fragments detected by Posterior decoding (sensitivity) while detecting more true archaic sequence than Viterbi (specificity), making the new decoding results more reliable.

### Sampling from the posterior and hybrid decoding methods applied to real data

Previous studies have reported that East Asians possess, on average, ∼20% more Neanderthal ancestry than Europeans (Vernot and Akey 2014; Sankararaman et al. 2014; Coll Macià et al. 2021). Additionally, differences in the length distribution of archaic fragments among Eurasian populations are linked to variations in ancestral generation times across these groups, as shorter archaic fragments imply more rounds of recombination – thus, more generations – since the admixture event (Coll Macià et al. 2021).

We sought to quantify more precisely these observations on Neanderthal ancestry proportions and fragment lengths using the sampling approach in hmmix applied to real data (Methods). For these analyses, we selected one representative individual from each of three major Eurasian populations within the 1000 Genomes Project dataset: NA19078 (Japanese in Tokyo, Japan [JPT] with predominantly East Asian ancestry), NA20810 (Toscani in Italia [TSI] with predominantly European ancestry) and NA21130 (Gujarati Indians in Houston, Texas, USA [GIH] with predominantly South Asian ancestry).

Often, Neanderthal ancestry proportions are quantified using the direct f4 ratio method (Petr et al. 2019; Bergström et al. 2020) (Methods). Our hmmix-based estimates of Neanderthal introgression for all three decoding methods are consistent with the broad confidence intervals reported by f4 ratio. Among hmmix decodings, sampling from the posterior consistently yields estimates closest to the mean f4 ratio (Figure 6a, Methods). In line with patterns observed in simulation analyses, the sampling method identifies the highest amount of Neanderthal-derived sequences, followed by Posterior decoding and then Viterbi decoding (Figure 6a). Notably, sampling from the posterior reproduces the higher Neanderthal ancestry proportion previously observed in East Asians (1.59 %, 95% CI: 1.58%–1.61%) compared to Europeans (1.18 %, 95% CI: 1.16%–1.19%)—a difference not significant using only the f4 ratio statistic, given its lower resolution with individual-level data.

**Figure 6.**
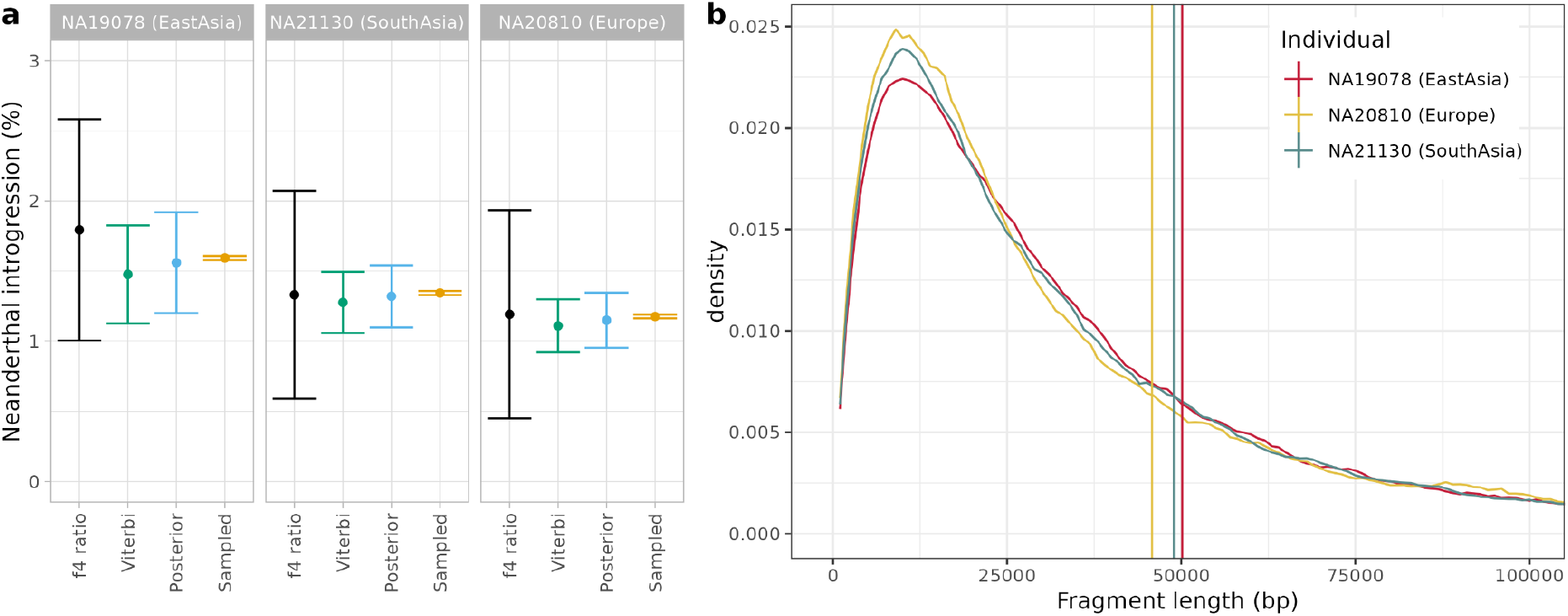
Archaic fragment summary statistics in three individuals a) Amount of Neanderthal ancestry in the phased genome for the three individuals using the f4 ratio statistic and three different hmmix decoding schemes. Error bars represent 95%CI of the mean for each estimate. For f4 ratio, the variance is estimated with jackknife with 0.050 cM block size (714 blocks). For Viterbi and Posterior decoding the variance is estimated with jackknife by chromosome. For the samples from the posterior, the 95%CI correspond to the 0.025 and 0.975 percentile values of the distribution sampled. b) Length distribution of introgressed archaic fragments for 3 individuals truncated at 100 kb. For each individual we performed 1,000 inhomogeneous samples. The vertical lines represent the mean fragment length for all 1,000 samples. Only fragments with at least one observation are used.

We also analyzed the samples from the posterior to estimate archaic fragment length distributions among the selected individuals (Figure 6b). While the mean fragments length is overestimated due to recombination not being uniform along the genome (Supplementary Material 4) we can still investigate the difference between individuals assuming the overall recombination map is similar across populations. Average fragment lengths are 45.81 kb (95% CI: 45.78–45.84 kb) for the European individual, 48.95 kb (95% CI: 48.92–48.98 kb) for the South Asian individual, and 50.14 kb (95% CI: 50.11–50.17 kb) for the East Asian individual. These fragment lengths are shorter than those previously reported by Coll Macià et al.(Coll Macià et al. 2021), likely due to our analysis using phased rather than diploid data and the greater sensitivity of sampling from the posterior for detecting shorter fragments compared to the Posterior decoding used in that study. Nevertheless, assuming reasonable parameters for recombination rates and admixture timing, our results remain consistent with previously inferred generation time differences of approximately 1 to 3 years among Eurasian populations since their admixture event with Neanderthals (Methods).

Next, we apply hybrid decoding to the three individuals and compare the identified archaic fragments with those obtained previously using Viterbi and Posterior decoding. We observe a clear nested relationship among the three methods: Posterior decoding consistently identifies the most fragments, encompassing all fragments found by hybrid and Viterbi decoding, while hybrid decoding similarly includes all fragments detected by Viterbi in all haplotypes of the three individuals (Figure 7a, Table S3). A small proportion (<5%) of fragments called by Viterbi decoding are subdivided into two or three smaller fragments by hybrid and/or Posterior decoding, indicating the more local resolution of these methods. Likewise, hybrid fragments were occasionally (<1.5%) further subdivided by Posterior decoding (Figure 7b, Table S3). Figure 7c provides an example of such cases in which the boundaries of archaic fragments identified by the three decoding methods differ significantly. The example involves a 126 kb genomic region on chromosome 1 of individual NA19078. This region contains two distinct clusters of variants shared with Neanderthals, upstream and downstream of 159.25 Mb. Viterbi decoding identifies only the second cluster as archaic, whereas Posterior decoding identifies two separate archaic fragments corresponding to each cluster individually. In contrast, hybrid decoding infers a single archaic fragment spanning both. Given the high linkage disequilibrium (R^2^ ∼0.8, Methods, Table S4) observed among the Neanderthal-shared variants in the flanking regions of these two clusters, the scenario suggested by hybrid decoding appears more plausible. This scenario suggests that both clusters represent parts of a single Neanderthal haplotype in contrast to Viterbi decoding which only identifies a fragment localized in the second cluster. Moreover, the Posterior decoding scenario with two fragments implies multiple recombination events within this limited genomic region, making it a less parsimonious explanation due to the high LD observed.

**Figure 7.**
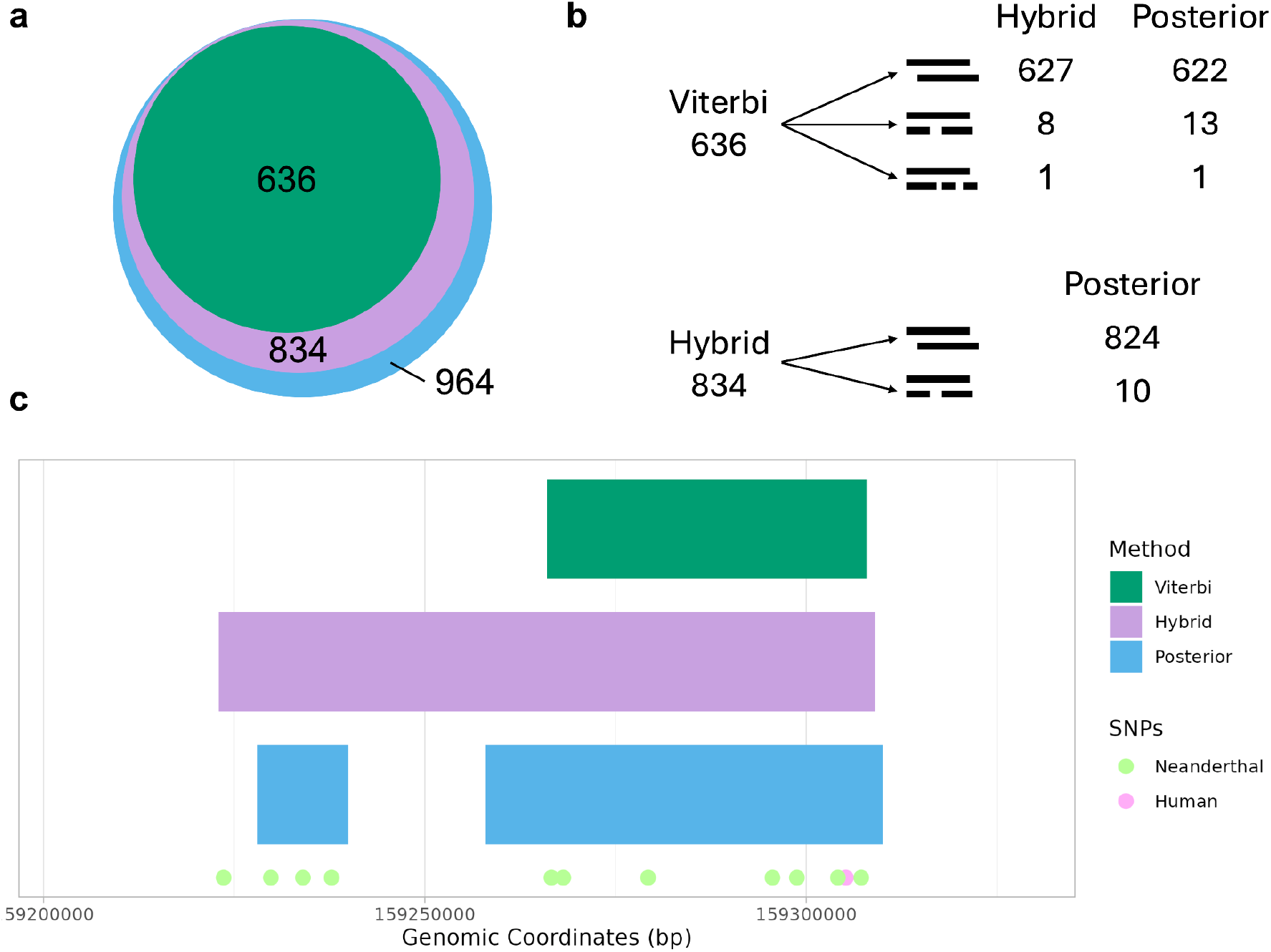
Hybrid decoding for individual NA19078, haplotype 1, chromosome 1. a) Venn diagram of the archaic fragments found by each decoding method. b) Number of fragments from Viterbi (above) and hybrid (below) that are found by a single (one to one fragment) or multiple fragments (one to two or one to three fragments) by the methods in the columns (hybrid and/or Posterior decoding). c) Local archaic fragment calling for a region in which there are differences in calling among methods. Decoded fragments for each method are shown as horizontal bars. Observed genetic variants are shown as circles. Variants shared with any of the high coverage Neanderthal genome or Denisova genome are painted accordingly. Variants not found in any archaic genome are painted as Human.

## Discussion

In this manuscript, we introduce two novel enhancements to hmmix which improve the detection of archaic fragments and estimation of summary statistics relevant in the study of the admixture events between modern and archaic humans. An updated version of hmmix incorporating these enhancements is available on https://github.com/LauritsSkov/Introgression-detection.

The first enhancement focuses on improving the estimation of summary statistics with an analytical and empirical technique based on conditioning the hidden state sequence distribution on the observed data. FMCI provides exact analytical distributions for summary statistics but becomes computationally intensive at genome-wide scales. Calculating archaic fragment length distribution or the longest fragment distribution can become infeasible when analyzing datasets as in this manuscript containing millions of observations. To overcome these computational challenges, we have divided and parallelized computations. However, future research should investigate more efficient strategies, such as adaptive matrix sizing or sparse matrix approaches. As for now, the dimensions of FMCI matrices must be pre-set based on parameters typically approximated with sampling from the posterior—an approach that introduces some circularity, given that these empirical summaries already serve this purpose.

In general, posterior sampling is computationally more efficient and the same hidden state samples simultaneously provide all summary statistics. We have shown how this approach significantly improves precision, particularly in estimating the proportion of archaic ancestry with individual genomes. Sampling from the posterior captures shorter and less probable fragments among all samples, which are never represented in Viterbi nor Posterior decoding, due to the lack of probabilistic support. Sampling from the posterior substantially reduces bias, specifically in fragment length summaries estimation. With the correct fragment length distribution it will enable more accurate and direct inference of admixture times with archaic populations in future studies. Finally, estimates derived from these approaches demonstrate robustness to violations of classical HMM assumptions, such as non-geometric distributions of fragment lengths, which can occur due to variable recombination rates or recent admixture events (Liang and Nielsen 2014).

The second enhancement, hybrid decoding, effectively combines the advantages of Posterior decoding and Viterbi. By adjusting the *α* parameter, researchers can smoothly transition between local (Posterior decoding) and global (Viterbi) decoding strategies, enabling optimal customization to specific analytical needs. The Artemis plot reveals that hybrid decoding successfully combines global decoding with local probabilistic precision, significantly reducing both false-positive and false-negative fragment identifications. This flexibility is particularly advantageous in targeted genomic analyses, facilitating informed assessments of archaic ancestry in particular haplotypes through decoding strategies positioned between the traditional Posterior decoding and Viterbi solutions.

Although specifically developed for hmmix, these methods can be adapted to any HMM framework. We anticipate these methodological enhancements will become widely adopted analytical tools, benefiting various HMM-based genomic studies and other fields.

## Methods

### Simulations

We introduce two distinct scenarios, each characterized by specific assumptions that constrain the parameters of the two-state Poisson HMM in hmmix (Figure 1): emission probabilities (Φ), starting probabilities (π), and transition probabilities (Γ).

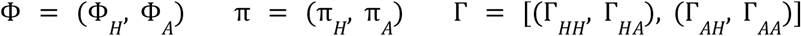

Where *H* denotes the human state and *A* denotes the archaic state. Note that Φ is determined by the Poisson rates *λ* = (*λ* _*H*_ , *λ* _*A*_) in the case of hmmix. From these HMMs data is simulated and analyzed.

#### 1) Realistic parameters

The realistic set is calculated such that the mean archaic fragment length is 50 kb and the archaic proportion genome wide is 2%.

The first assumption defines the transition probability between archaic states (Γ _*AA*_ ), since the archaic fragments are geometrically distributed with rate 1 − Γ _*AA*_. Thus, the mean of that distribution is

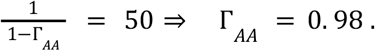

Additionally, we have that

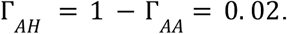

The second assumption defines the starting parameters π_*A*_ = 0. 02 and π _*H*_ = 0. 98. Since the starting probabilities (π) are the stationary distribution of the Markov chain, it holds that

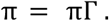

Together with Γ_*AA*_ , we obtain the remaining parameters with

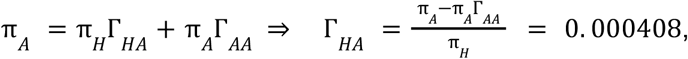

and

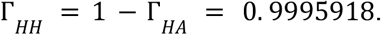

Finally, the emission parameters – which depend on the divergence time between modern humans and archaics (archaic state emissions) and the divergence time between African and nonAfrican genomes (human state emissions) – are obtained from literature (Skov et al. 2020) and set to *λ*_*A*_ = 0. 3 and *λ*_*H*_ = 0. 03.

These parameters are presented in Table 1 and saved in the file `realistic_param.json`.

#### 2) Visualization parameters

The second set of parameters increases the overall amount of archaic sequence (2/3) and shortens the length of human fragments (mean length = 100 kb). We also make the emission probabilities more similar between the two states so that decoding methods have less power to detect archaic fragments.

This way, we obtain more archaic fragments overall and more differences among decoding methods for visualization purposes. These parameters are presented in Table 3 and saved in the file `visualization_param.json`.

**Table 3.** Visualization parameters and their Baum-Welch estimated parameters.

| $a$ ($\times 10^4$ years) | $r$ ($\times 10^{-8}$ bp x generation) | $\bar{L}_{EUR}$ ($\times 10^3$ bp) | $\bar{L}_{EAS}$ ($\times 10^3$ bp) | $\Delta_{GT}$ (years) |
| --- | --- | --- | --- | --- |
| 6.00 | 0.80 | 45.81 | 50.14 | 2.08 |
| 4.00 | 0.80 | 45.81 | 50.14 | 1.39 |
| 6.00 | 1.25 | 45.81 | 50.14 | 3.25 |
| 4.00 | 1.25 | 45.81 | 50.14 | 2.17 |

Simulated data of a single chromosome of 1×10^6^ 1kb-windows (genomic sequence of 1 Gb) are generated with the hmmix command `hmmix make_test_data` for both realistic and visualization parameters.

Parameters are estimated with the implemented Baum-Welch algorithm in hmmix by running the command `hmmix train` (Table 1 and Table 2). We use the estimated parameters for all the analyses in the manuscript.

The 95% CI for parameter estimation is obtained by bootstrapping. For that, we simulate data from the parameter estimates (Table 1) and re-estimating parameters using the Baum-Welch algorithm 100 times with hmmix (`hmmix make_test_data` and `hmmix train` commands). The interval is obtained for each parameter as the 2.5% and 97.5% percentile of the 100 parameter values estimated. The 95% CIs are very narrow, with little variation and capture the true value (considering up to 4 decimal digits). This reflects that there is little uncertainty in the parameter estimation in the simulated data by hmmix.

The exact commands and relevant files are provided in the simulated data and the real data directories on the GitHub repository (https://github.com/MoiColl/HMMenhancements).

### Sampling from the posterior

As shown in Bæk et al. 2025, the hidden state sequence *y* conditional on the observed sequence *x* is an inhomogeneous Markov chain with transition probabilities

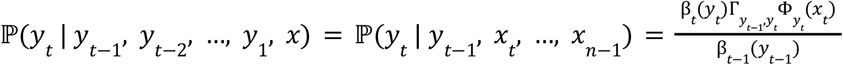

for *t* = 1, …, *n* − 1 where β _*t*_ (*y* _*t*_ ) = ℙ(*x* _*t*+1_ , …, *x* _*n*−1_ | *y*_*t*_ ) is the matrix of backward probabilities from the Posterior decoding framework, Γ is the matrix of transition probabilities and Φ is the emission probabilities. In the case of hmmix, the emission probabilities Φ correspond to the Poisson rates *λ*, such that Φ_*A*_ (*x*_*t*_ ) = *Poisson*(*x*_*t*_ ; *λ*_*A*_ ) and Φ _*H*_ (*x* _*t*_ ) = *Poisson*(*x*_*t*_ ; *λ*_*H*_ ), which are the probabilities of observing *x* SNPs in position *t* under a Poisson distribution in the archaic state *A* and the human state *H*, respectively.

The initial distribution for *t* = 0 of the inhomogeneous Markov chain is

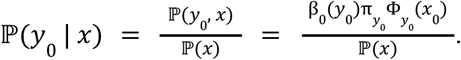

See Bæk et al. 2025 for the proof of the theorem above. Note that here the first index is 0 while in Bæk et al. 2025 the first index is 1. This is because the code provided in this manuscript is written in python.

The probabilities of staying in the human state conditional on the data ℙ( *y* _*t*_ = *H* | *y* _*t*−1_ = *H, x*_*t*_ , …, *x* _*n*−1_ ) are denoted throughout the paper as *a* and the probability of leaving the human state ℙ( *y*_*t*_ = *A* | *y*_*t*−1_ = *H, x*_*t*_ , …, *x*_*n*−1_ ) as 1 − *a*_*t*_. Similarly, the probabilities of staying in the archaic state conditional on the data ℙ( *y*_*t*_ = *A* | *y*_*t*−1_ = *A, x*_*t*_ , …, *x*_*n*−1_) are denoted as *b*_*t*_ and the probability of leaving the archaic state ℙ( *y*_*t*_ = *H* | *y*_*t*−1_ = *A, x*_*t*_ , …, *x*_*n*−1_) as 1 − *b*_*t*_. Similarly denoted, the initial values ℙ(*y*_0_ = *H* | *x* ) are denoted *a*_0_ and ℙ(*y*_0_ = *A* | *x* ) as *b*_0_ .

Once the HMM parameters have been estimated and given the observed sequence, these inhomogeneous transition probabilities can be used to sample hidden state sequences.

The calculation of these inhomogeneous transition probabilities, as well as sampling hidden state sequences, are implemented in hmmix with the command `hmmix inhomogeneous`. In this study, 1,000 sequences are sampled for both realistic and visualization simulated scenarios and for each real data individual.

The exact commands are provided in the simulated data and the real data directories on the GitHub repository (https://github.com/MoiColl/HMMenhancements).

### Finite Markov Chain Imbedding (FMCI)

The FMCI framework consists on building a Markov chain for a specific pattern of interest at every point in the sequence based on the inhomogeneous transition probabilities *a*_*t*_ and *b*_*t*_. Once the chain is encoded in a matrix Δ(*a*_*t*_, *b*_*t*_) for every point in the sequence (*t*), we can obtain the probability distribution of the summary of interest with

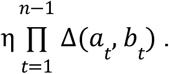

Here *η* represents the starting vector consisting of *a*_0_ and *b*_0_ . The resulting vector will encode the probability distribution of the summary of interest for the sequence analyzed.

The number of archaic fragments as an example is explained in detail in Supplementary Material 1. The FMCI constructed to compute the number of archaic fragments (number of jumps), the total archaic sequence (number of states), the maximum archaic fragment length (longest run) and the archaic fragment length distribution (run length distribution) are detailed in Bæk et al. 2025. The running time for all algorithms as well as strategies to parallelize and decrease running time are explained in Supplementary Material 1.

The FMCI scripts are provided in the FMCI directory on the GitHub repository (https://github.com/MoiColl/HMMenhancements).

### Hybrid decoding

Viterbi decoding inferes the global hidden state sequence *s* that maximizes ℙ(*y* = *s* | *x* ), where *x* is the observed sequence and *y* the hidden state sequence

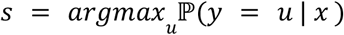

Posterior decoding inferes the local hidden state sequence *s*_*t*_ that maximizes ℙ(*y*_*t*_ = *s*_*t*_ | *x* ) for every position *t* in the sequence

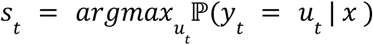

The hybrid decoding method mixes Posterior decoding and Viterbi methods by combining the logarithm function both in the same equation as

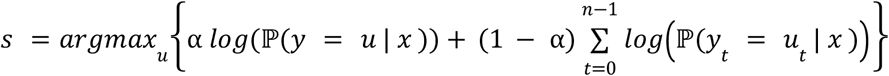

where *α* ∈ [0, 1] weights the two terms in the equation resulting in the same solution as Posterior decoding when *α* = 0, the same solution as Viterbi decoding when *α* = 1 and intermediate values correspond to hybrid solutions.

The *α* parameter can be chosen using the Artemis analysis as described in the main text of this paper and Bæk et al. 2025, and the hybrid path for a given *α* can be calculated efficiently using a recursive algorithm similar to Viterbi (Kuljus and Lember 2023, Bæk et al. 2025). The full overview of the method and its derivation can be found in Bæk et al. 2025.

The hybrid decoding method is implemented in hmmix using the command `hmmix decode -hybrid`. The pipeline to find the optimal *α* using Artemis analysis is also implemented in hmmix with the command `hmmix artemis`. The exact commands are provided in the simulated data’s jupyter notebook and the real data’s `workflow.py` on the GitHub repository (https://github.com/MoiColl/HMMenhancements).

### Real Data Sources

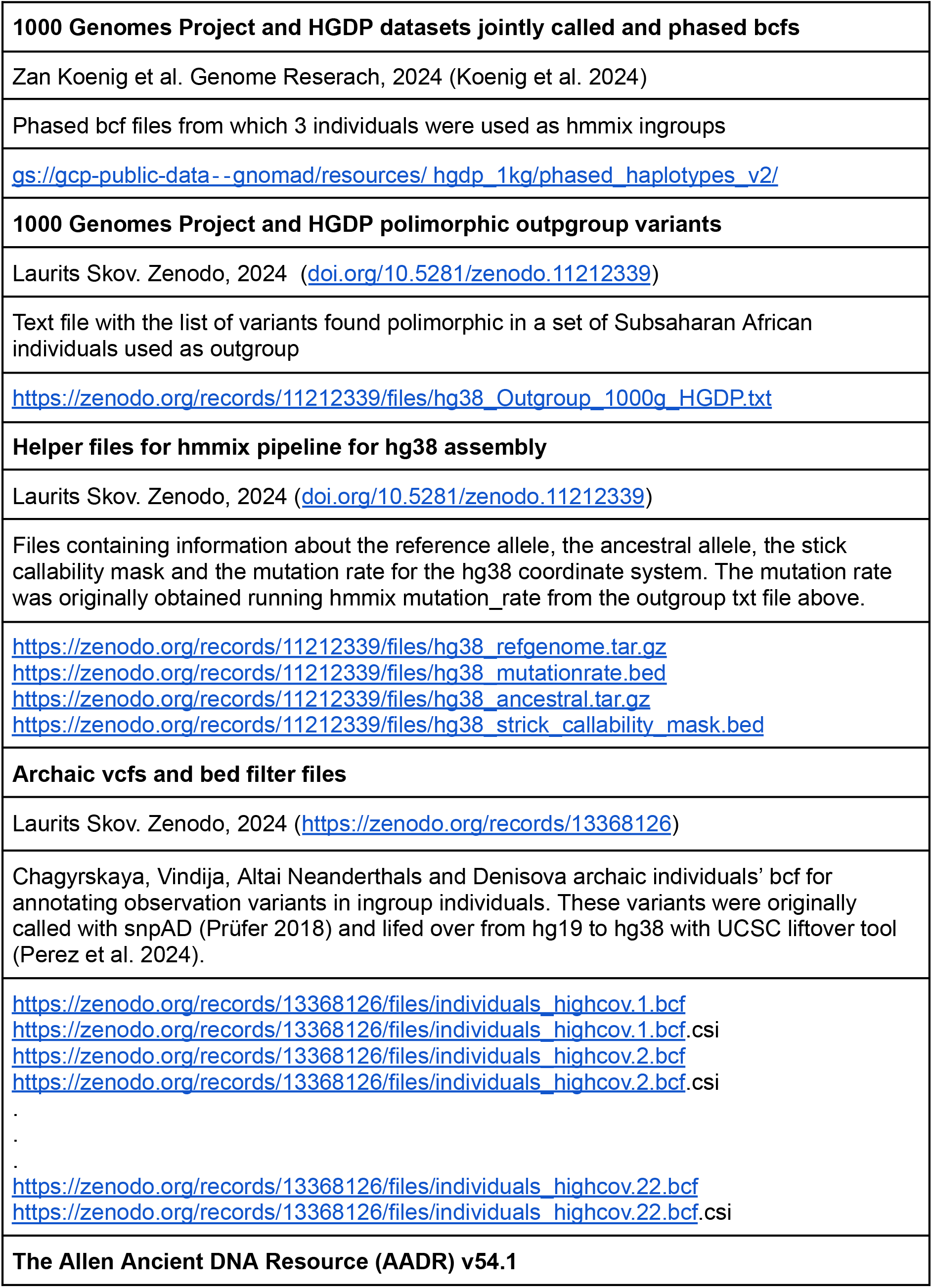
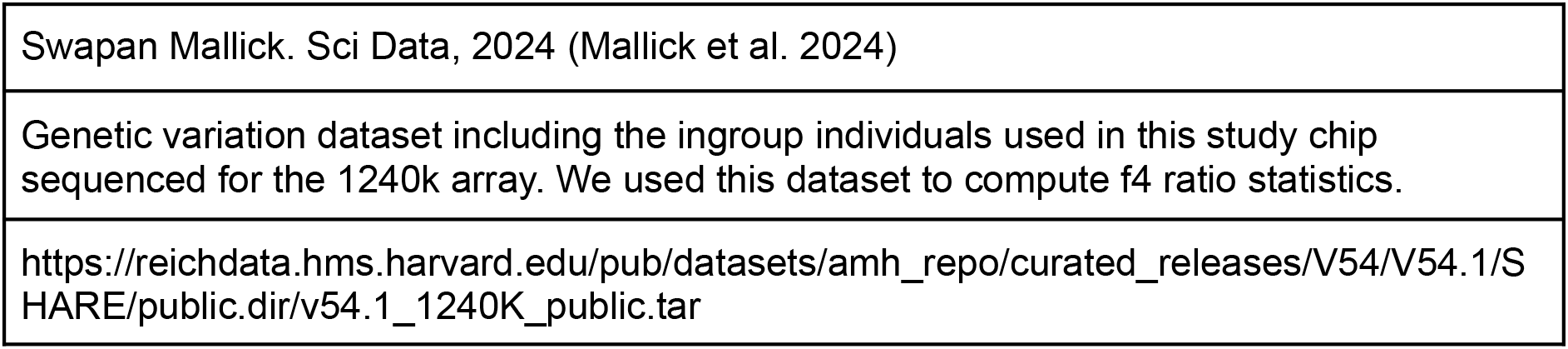

### Direct f4 ratio statistic

To compute the f4 ratio we used qpF4ratio (https://github.com/DReichLab/AdmixTools). We used the AADR dataset.

We run the statistic

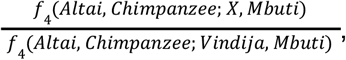

where *X* is any of the three test individuals NA19078, NA20810 and NA21130. The exact commands are given in the real data’s jupyter notebook on the GitHub repository (https://github.com/MoiColl/HMMenhancements).

### Historical generation time comparison among individuals

The archaic fragment length distribution (*L*) of a given individual is exponentially distributed and the rate of decay (*λ*) corresponds to the number of generations since the admixture event (*g*) and the recombination rate per generation (*r*), thus

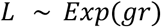

Since the mean of an exponential distribution is the inverse of its rate, we have that the mean archaic fragment length can be calculated as

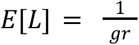

We can rewrite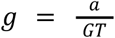, where *a* is the years since the admixture event and *GT* is generation time. Thus, given *E*[*L*], *a*, and *r*, we can get *GT* with

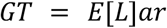

The expected fragment length can be approximated with the average fragment length 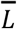

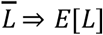

To know the difference in generation time between population *X* and population *Y*, we can compute that with

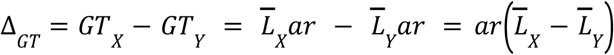

If we assume reasonable values for *a* and *r*, and use the mean fragment length from the European individual (NA20810) and the East Asian individual (NA19078), we get

Thus a difference of 1 to 3 years of generation time since the admixture event.

### Linkage disequilibrium patterns

We calculated r2 with vcftools (v 0.1.16) (Perez et al. 2024) among East Asian individuals from the real data bcfs. The exact commands are provided in the real data’s jupyter notebook on the GitHub repository (https://github.com/MoiColl/HMMenhancements).

## Code availability

The updated hmmix 1.0.0 can be installed through pipy and can be found in https://pypi.org/project/hmmix/ and https://github.com/LauritsSkov/Introgression-detection.

The scripts coded to produce data and tables, perform statistical analysis and plot figures for this manuscript are accessible on GitHub (https://github.com/MoiColl/HMMenhancements). The scripts provided in this repository are licensed under the MIT License.

## Acknowledgements

All of the computing for this project was performed on the GenomeDK cluster. We would like to thank GenomeDK and Aarhus University for providing computational resources and support.

## Supplementary Material

### S1 Finite Markov Chain Imbedding (FMCI)

The FMCI framework consists of building a Markov chain to compute a summary statistic of interest based on the inhomogeneous transition probabilities. Below, we explain the details of computing FMCI for the number of archaic fragments as a general illustrative example. The description of the other summary statistics can be found in Bæk et al. 2025. Here, we only discuss the running time of the algorithms corresponding to each statistic and how to speed up those by mainly parallelizing computations.

#### Number of archaic fragments

Taking as example the calculation of the number of archaic fragments in a sequence, the corresponding states of the Markov chain can be coded with two indices (Figure S1):

1. the number of archaic fragments observed (first index, starting at 0 with a maximum number of *l* fragments)
2. being in an archaic or human state (second index, archaic “A” or human “H”)

Considering this, there will be 2*l* + 2 states, plus a final absorbing state, thus a total of 2*l* + 3 states defining the Markov chain (Figure S1). The first two states (index 0 and 1), correspond to the probabilities of observing 0 archaic fragments. The following two states (index 2 and 3), correspond to the probabilities of observing 1 archaic fragment and so on. Finally, the absorbing state will hold the probability of observing *> l* fragments.

The transition probabilities of the defined Markov chain can be represented with a square matrix Δ(*a*_*t*_, *b*_*t*_) in which appear the probabilities of transitioning from one state to another for a specific *t* in the sequence. More specifically, the row index correspond to the state that transition start from (*t* − 1), and the columns correspond to the transition end (*t*, Figure 7b).

**Figure S1.**
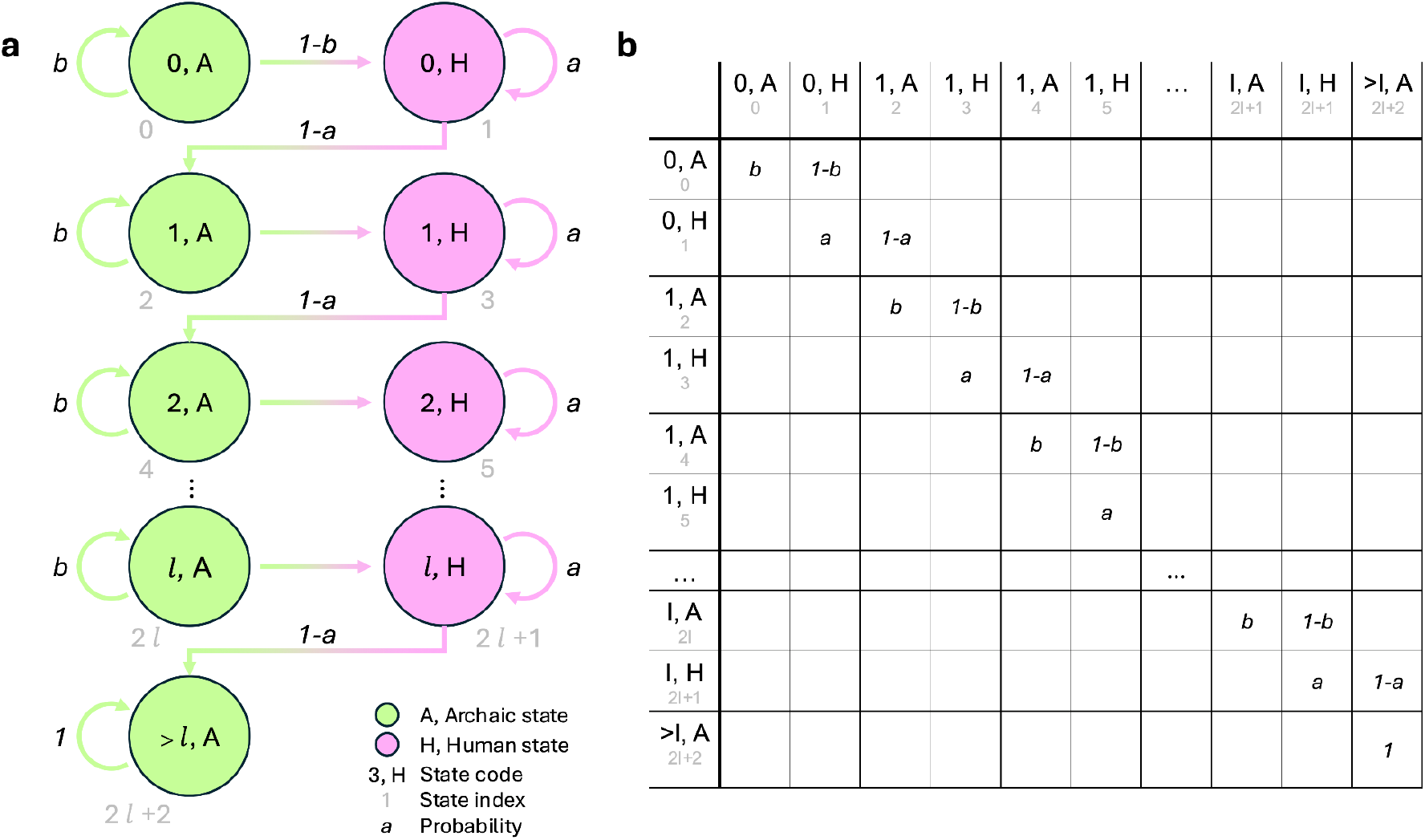
Markov chain Δ(*a*_*t*_ , *b*_*t*_ ) representation of the FMCI method in graph form (a) and its transition probability matrix (b) to compute the number of archaic fragments in a sequence. This figure is based on figures and matrices from Bæk et al. 2025, adapted to the hmmix case. Note that in Bæk et al. 2025, states H and L denote High and Low respectively, and do not relate to the states used here, Human (H) and Archaic (A). Also note that state index is a 0-start-index, while the corresponding index in the Bæk et al. 2025 is 1-start-index.

Note that the transition probabilities *a* and *b* are time dependent, meaning that they change depending on the position *t* in the sequence of length *n* that is analyzed.

The initializing probabilities (*t* = 0) to compute the number of archaic fragments is *η* = (0, *a*_0_ , *b*_0_ , 0, 0, …, 0 ). The first entry is defined as 0 since according to the transition matrix, that state would correspond to start in an archaic fragment, but counting 0 fragments. Thus, if starting in an archaic fragment, the count should directly increase to 1, which corresponds to state 2.

The posterior distribution for the number archaic fragments for the entire sequence can be calculated from

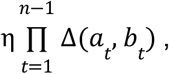

which results in a vector of length 2*l* + 3. The sum of the two first entries corresponds to the probability of zero fragments, the sum of the next two entries is the probability of one fragment, and so on. The last entry corresponds to the probability of observing *l* + 1 or more fragments.

Thus, the time complexity of this algorithm will be *O*(*nl*^2^). For simulations, *n* is 10^6^. To reduce the computational time of the algorithm, we minimize *l* by approximating this value with the results from sampling from the posterior, which we set to 10^4^ (Figure 3).

#### Total archaic sequence

The running time for the total archaic sequence is

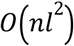

Where *n* is the number of states in the sequence (genome length, 10^6^ for simulations), and *l* is the maximum archaic sequence that the probability will be computed for which can be approximated with the samples from the posterior (∼2.05×10^4^ for simulations).

Since *l* is large, the algorithm’s running time becomes computational infeasible to complete. Thus, we use the strategy of dividing the sequence in 100 blocks of size 10^4^ to be analyzed independently. This enables us to reduce both *n* to 10^4^ and *l* to 10^3^, making each computation much faster. The results are then convoluted to sum all the distributions.

Note that the last probability in each block corresponds to observing *> l* archaic sequence. When performing convolution, operations involving this last bin incorrectly attribute probability mass to specific outcomes, even though it actually represents an aggregate of larger, unbounded values. As a result, the probability mass in the convolved distribution may be slightly inflated. Since we choose *l* large enough the last bin has very little probability mass, the effect on the resulting distribution is negligible. This is also true for the archaic fragment length distribution, for which we also convolve.

#### Archaic fragment length distribution

This algorithm computes the probability distribution of observing the number of archaic fragments of extract length *k*. Once we get the probability distribution, it reports the mean number of fragments of extract length *k*. To obtain the full distribution of lengths 1, 2, 3, …, *K* we must compute the mean value for every value *k* independently. The time complexity for each computation is

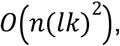

where *n* is the number of states in the sequence (genome length, 10^6^ for simulations), *k* is the fragment length (up to 250), and *l* is the maximum number of archaic fragments of exact length *k* for which we are going to compute the probability for. Similarly to the total archaic sequence, we reduce *n* by dividing the sequence in 2 chunks to be analyzed independently and the resulting distributions are subsequently convoluted. Based on the results from sampling from the posterior distribution, we also limit *l* to 25 for *k* ≤ 50, to 15 for 25 < *k* ≤ 100, to 10 for 100 < *k* ≤ 150, to 5 for 150 < *k*, which is critical for the running time of the algorithm. Running times for all computations are < 1 hour, with a big majority being < 30 min.

#### Longest archaic fragment

The time complexity of this algorithm is

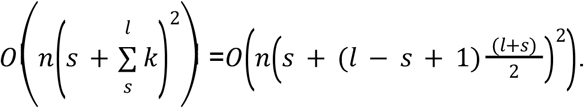

In the worst case scenario *s* = 1 we get

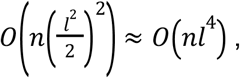

where *n* is the number of states in the sequence (genome length, 10^6^ for simulations), *s* and *l* are the minimum (260 in simulations) and maximum (380 in simulations) archaic fragment length respectively for which a probability will be computed and for all lengths in between. The first strategy to reduce computational time is to minimize the distance between *s* and *l*. We set *s* to 260 and *l* to 380 by approximating the corresponding values from the samples from the posterior (Figure 3). The second strategy is to compute sections of the distribution independently. For example, if we compute sections of length *z*, the first section would compute from *s* to *s* + *z*, and the second *s* + *z* to *s* + 2*z*, and so on. The resulting sections can be aggregated together to form the probability distribution *s* to *l*. We set *z* to 5 in the simulation study. Running times for all computations are < 4 hours.

### S2 Robustness to model misspecifications

We test the robustness of the sampling from the posterior approach to misspecification and violations of the following HMM assumptions:

1. When there is very little differentiation of the emission parameters from the two states. This would correspond to coalescent times between Africans and Non-Africans being closer to coalescent times between archaic and modern humans.
2. When the number of observations does not follow a Poisson distribution. This corresponds to coalescent events between modern humans and archaics not happening at the same time across the genome, but having some variation.
3. When the archaic fragment length distribution does not follow a geometric distribution. This corresponds to when, for example, recombination is not uniform along the genome.

A helpful analogy to the inhomogeneous sampling process is Bayesian statistics. One can think of the set of parameters π, Γ and Φ as the priors, the observations corresponds to the data, and sampling approach helps us obtain the posterior distribution.

When there is no signal in the data we would expect the posterior distribution to resemble the prior. When the signal in the data is very strong (e.g. when the emission probabilities are very different) the prior has little impact on the posterior.

We focus on the ability to capture the true summary statistics such as fragment length distribution, number of fragments, total sequence, accuracy and precision. Testing the sampling approach will, by extension, also serve as a test of the FMCI.

#### Differentiation of emission parameters

We simulate data with the realistic parameters (Table 1).We vary *λ*_*A*_ from the original realistic parameters to the values 0.031, 0.06, 0.09, 0.12, 0.15, 0.3, 1, 2, 3 to evaluate how robust the inhomogeneous method is to the difference between the emission parameters in the two states.

We simulate 10^6^ windows of sequence. For each simulated dataset we perform 100 inhomogeneous samples with both the known parameters (the parameters which generated the data) and the estimated parameters obtained from Baum Welch training. We show the results in Figure S2.

When we provide the inhomogeneous sampling approach with the known parameters we obtain distributions of the summary statistics which overlap the true values no matter how differentiated the emission parameters *λ*_*A*_ and *λ*_*H*_ are. This is due to the fact that we are always providing the correct prior even when there is no signal in the data. While the summary statistics are well captured the accuracy and precision are very low.

In general the estimated *λ*_*A*_ are within 2% of the true parameter value. The exceptions are when *λ*_*A*_ is 0.031 and 0.06 (very similar to *λ*_*H*_ which is 0.03). Here the trained *λ*_*A*_ are 0.0547 and 0.0542 respectively. The incorrect estimation of parameters means we have an incorrect prior. Since there is little signal in the data, this leads to incorrectly estimated fragments and thus summary statistics.

However, for the case of archaic admixture in modern humans, we are in a situation in which the Poisson rate from the archaic hidden state is ∼ 0. 3, and for that regime, sampling from the posterior shows robustness to recover the right summary statistics (Skov et al. 2018; Skov et al. 2020). In this regime, the sampling approach is also superior in capturing the summary statistics compared to the Viterbi and Posterior decoding.

**Figure S2.**
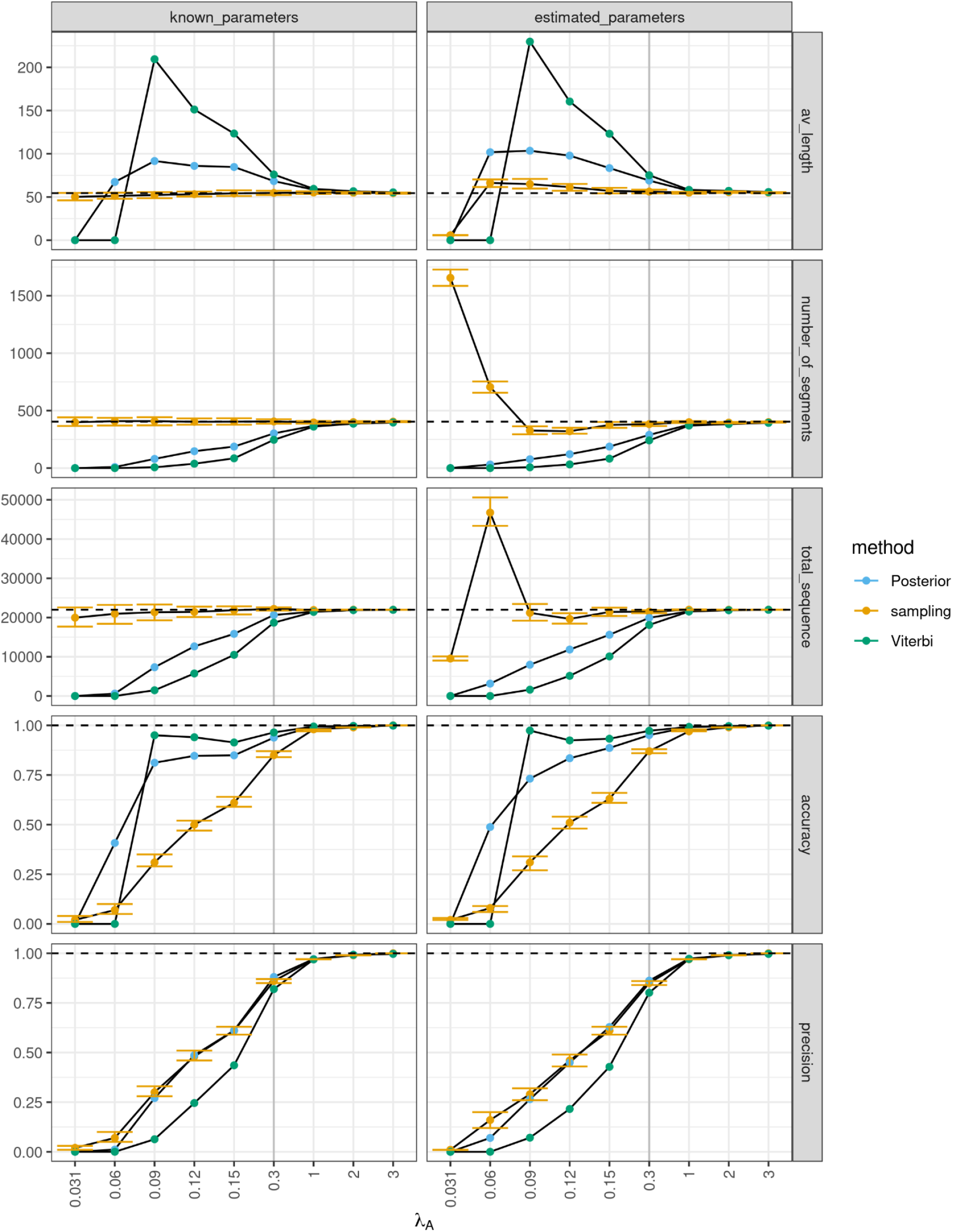
The dotted line indicates the true simulated values for each summary statistic shown in horizontal panels. We show the effect of varying *λ*_*A*_ for 100 samples where we use the true parameters (left panels) and the trained parameters (right panels). The grey vertical line shows the archaic emission parameter that is appropriate for studying archaic introgression in present-day humans.

#### Observation generated from an overdispersed Poisson process

hmmix assumes that observations are generated from a Poisson process - this corresponds to assuming that all fragments coalesce at a single time point. We now test the robustness of the sampling approach when the observations are generated from an overdispersed distribution.

For this test, we use the realistic parameters (Table 1), but now generate observations in both the human and archaic state from a negative binomial distribution where the mean is 0.03 and 0.3, respectively, but the variance changes. We express the overdispersion as the ratio of the variance to the mean.

The first simulation ratio of 1 which is equivalent to generating observations from a Poisson distribution. We next increase the ratio to 1.03125, 1.0625, 1.125, 1.25 and 1.5. We show the results in Figure S3.

We simulate 10^6^ windows of sequence. For each simulated dataset we obtain 100 samples from the posterior with both the known parameters (the parameters which generated the data) and the estimated parameters obtained from Baum Welch training.

The sampling approach fails to obtain the true summary statistics when the variance/mean ratio is greater than 1.0625. This is true whether or not we use the known or estimated parameters. This can be explained by the signal in the data (the overdispersion) having a larger impact than the prior.

**Figure S3.**
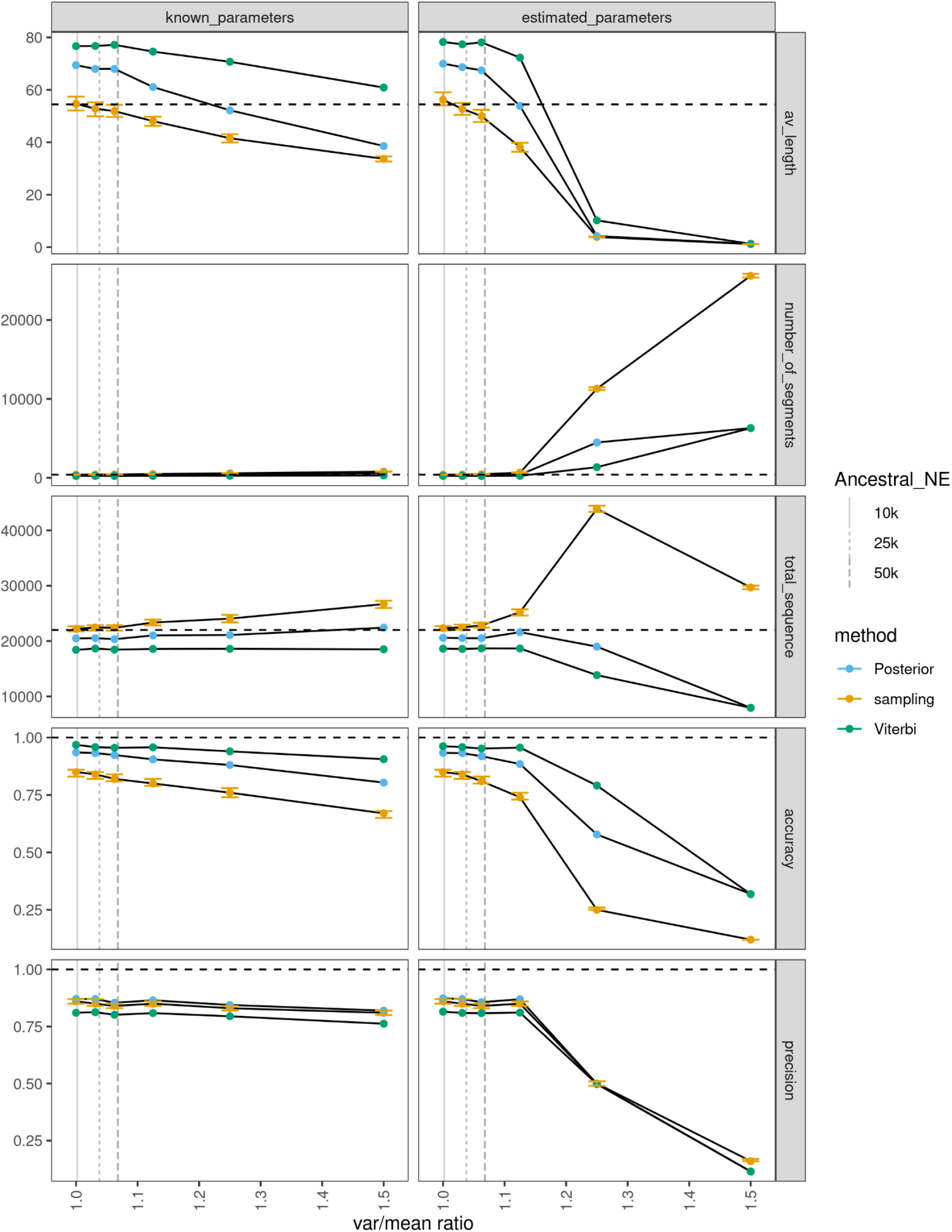
The dotted horizontal line indicates the true simulated values for each summary statistic shown in horizontal panels. We show the effect of increasing the variance of the negative binomial function compared to the mean for 100 samples where we use the true parameters and the trained parameters.Vertical dotted lines show the var/mean ratio for simulations with different ancestral effective population sizes.

We investigate what ratio of the variance/mean can be expected when analysing present-day human genomes. Recall that the emission parameter is related to the minimum coalescence time with the outgroup. The minimum coalescence time with the outgroup follows an exponential distribution with a parameter that depends on the effective population size and number of samples from the outgroup. Thus increasing the effective population size of the ancestral population will increase the mean coalescence time. Since it’s an exponential distribution the variance will increase more than the mean - thus increasing the ratio of the variance divided by the mean.

We focus on the emission parameter for the human state (a similar argument can be made for the archaic state) and set the effective population size of the ancestral population of Africans and non-Africans to 1,000, 10,000, 25,000 and 50,000 individuals. We name these simulation scenarios 1-4 respectively (Table S1). For each scenario, we simulate a non-African genome and 500 African genomes from a simple demography with the following parameters: The non-African and African population split 50,000 years ago. The effective population size of Africans after the split is constant effective population size at 20,000 individuals. The non-African population goes through a bottleneck where the effective population size reduces to 1,000 individuals for in the time interval 45,000 - 50,000. After that it recovers to 10,000 individuals.

The yaml files along with scripts for simulating the scenarios are provided in the GitHub folder (https://github.com/MoiColl/HMMenhancements).

For each scenario, we simulate 10,000 independent 1 kb windows and count the number of variants which are not seen in the 500 African genomes. We calculate the average number of SNPs per 1 kb window, i.e. the emission parameter for the human state. We also calculate the ratio of the variance divided by the mean. We show the results in Table S1.

**Table S1.**
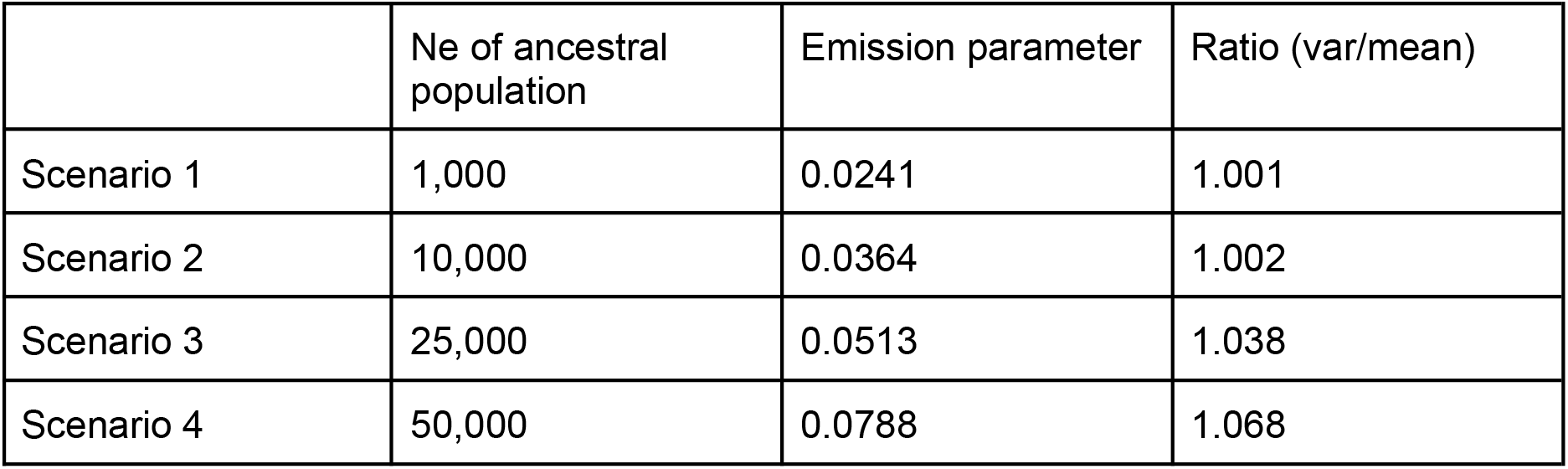
Simulation parameters for increasing variance to mean ratio of the emission rates.

It is often assumed that the effective population size of humans is below 20,000 (https://popsim-consortium.github.io/stdpopsim-docs/stable/catalog.html#sec_catalog_homsap_models_outofafricaextendedneandertaladmixturepulse_3i21)(Lauterbur et al. 2023).

If we assume an effective population size of non-African/African of 10,000 individuals we expect an emission parameter around 0.03 and a low degree of overdispersion (variance/mean ratio of 1.002) leading to accurate estimation of summary statistics with the sampling method. Even if the effective population size of non-African/Africans increases to 50,000 the summary statistics are well estimated. We show the variance/mean ratios for effective population sizes of 10k, 25k, 50k in Figure S3.

#### Fragment length distribution of archaic states generated from non-geometric distribution

Sampling from an homogeneous HMM produces a fragment length distribution that is geometric. In the absence of any signal in a data differentiating the two states, sampling from an inhomogeneous hmm will also produce a geometric distribution. We wanted to investigate under which circumstances the inhomogeneous sampling approach can recover non-geometric distributions.

The length distribution of a given hidden state (e.g, archaic state *A*) is geometrically distributed with the rate equal to the probability of leaving the state (e.g, 1 − Γ_*AA*_ ) under HMM assumptions. Here, we want to test if sampling from the posterior can capture length distributions that depart from the geometric assumption.

We first test the sampling approach to capture a combination of a Poisson distribution and a geometric distribution. We then proceed to test the performance of the sampling approach in the presence of a varying recombination rate which will tend to skew the distribution towards shorter fragments.

We simulate 10^6^ windows of data from the realistic parameters (Table 1) varying *λ*_*A*_ = 0.06, 0.3 and 3 and under two scenarios.

1. In scenario 1, 75% of archaic fragments follows a geometric distribution with parameter 1 − Γ_*AA*_ mean = 50. and 25% of archaic fragments follow a Poisson distribution with
2. In scenario 2, 100% of the archaic fragments are generated from a Poisson distribution with mean=50.

We show the results in Figure S4. When the emission parameters are very similar between the human and the archaic state (*λ*_*A*_ = 0.06 and *λ*_*H*_ = 0.03) there is not enough signal in the data to recover the true length distribution. The sampled length distribution is instead very similar to the geometric distribution of the estimated parameters. When the emission parameters become sufficiently distinct (*λ*_*A*_ ≥ 0.3 and *λ*_*H*_ = 0.03) the sampling approach approximates better the true length distribution.

**Figure S4.**
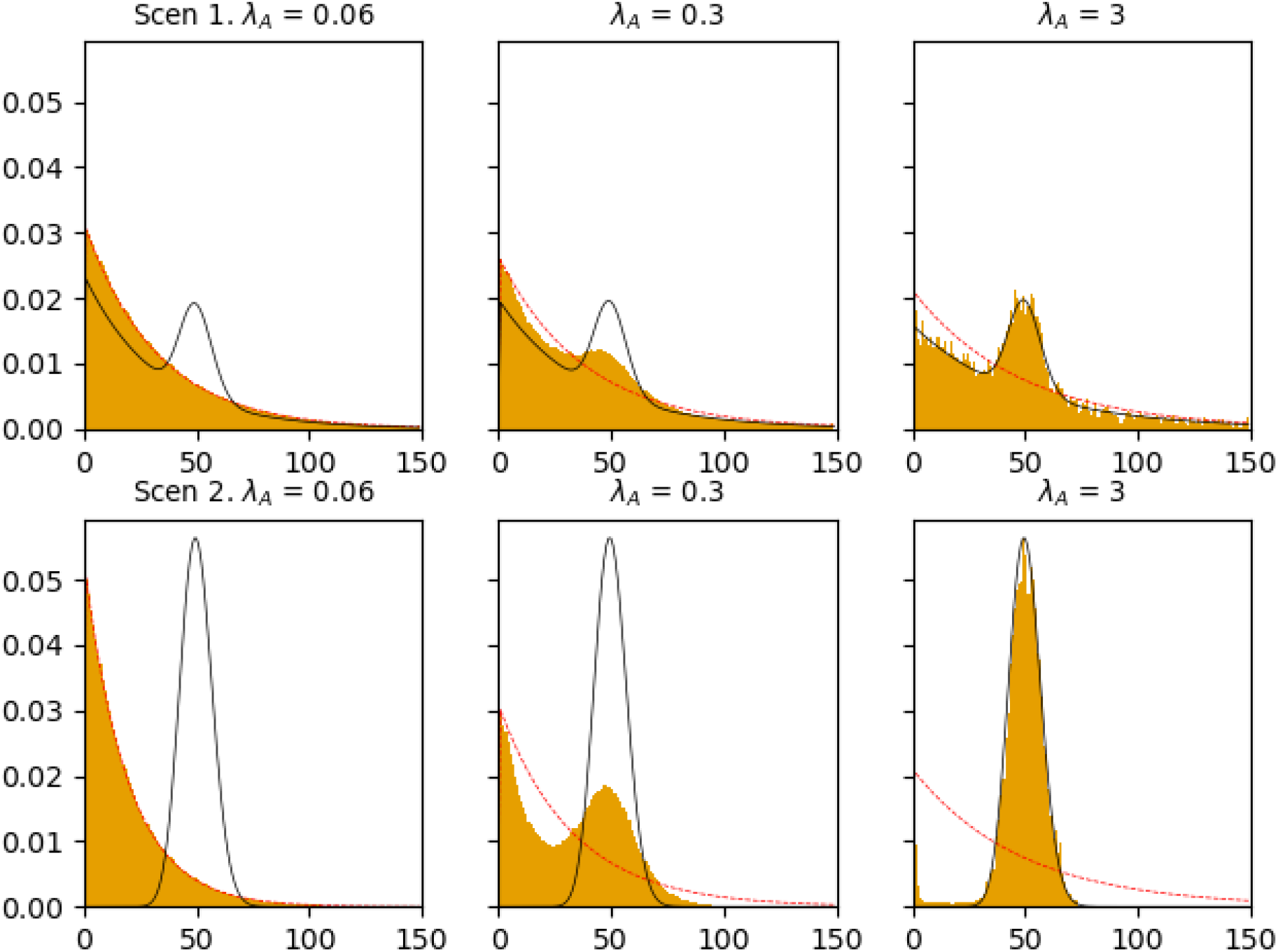
Length distribution from 100 samples from the data under two different scenarios and with three different emission values for the archaic state. The black line shows the true length distribution, the red line shows the geometric distribution with a parameter equal to the estimated transition parameter from the archaic state (1 − Γ_*AA*_ ), and the histogram is the length distribution from the 100 samples from the posterior.

We next investigate if the inhomogeneous sampling approach can capture the archaic fragment length that is generated by a constant and varying recombination rate across the genome respectively.

Using msprime (Baumdicker et al. 2021), we simulate a complete genome of 2.8 Gb with constant and varying recombination rate. We extracted true archaic introgressed fragments and added SNPs to the genome using *λ*_*H*_ = 0.03 for human fragments *λ*_*A*_ = 0.06, 0.3 and 3 for archaic fragments. We train the HMM parameters and decode the hidden state sequence using Viterbi and Posterior decoding algorithm. We also sample 100 paths from the conditional posterior probabilities.

For the simulations with constant recombination rate the fragment length distribution follows a geometric distribution and the sampling approach recovers the correct distribution even for *λ*_*A*_ = 0.06. For *λ*_*A*_ = 3, Viterbi, Posterior decoding and sampling approach all recover the correct fragment length distribution.

**Figure S5.**
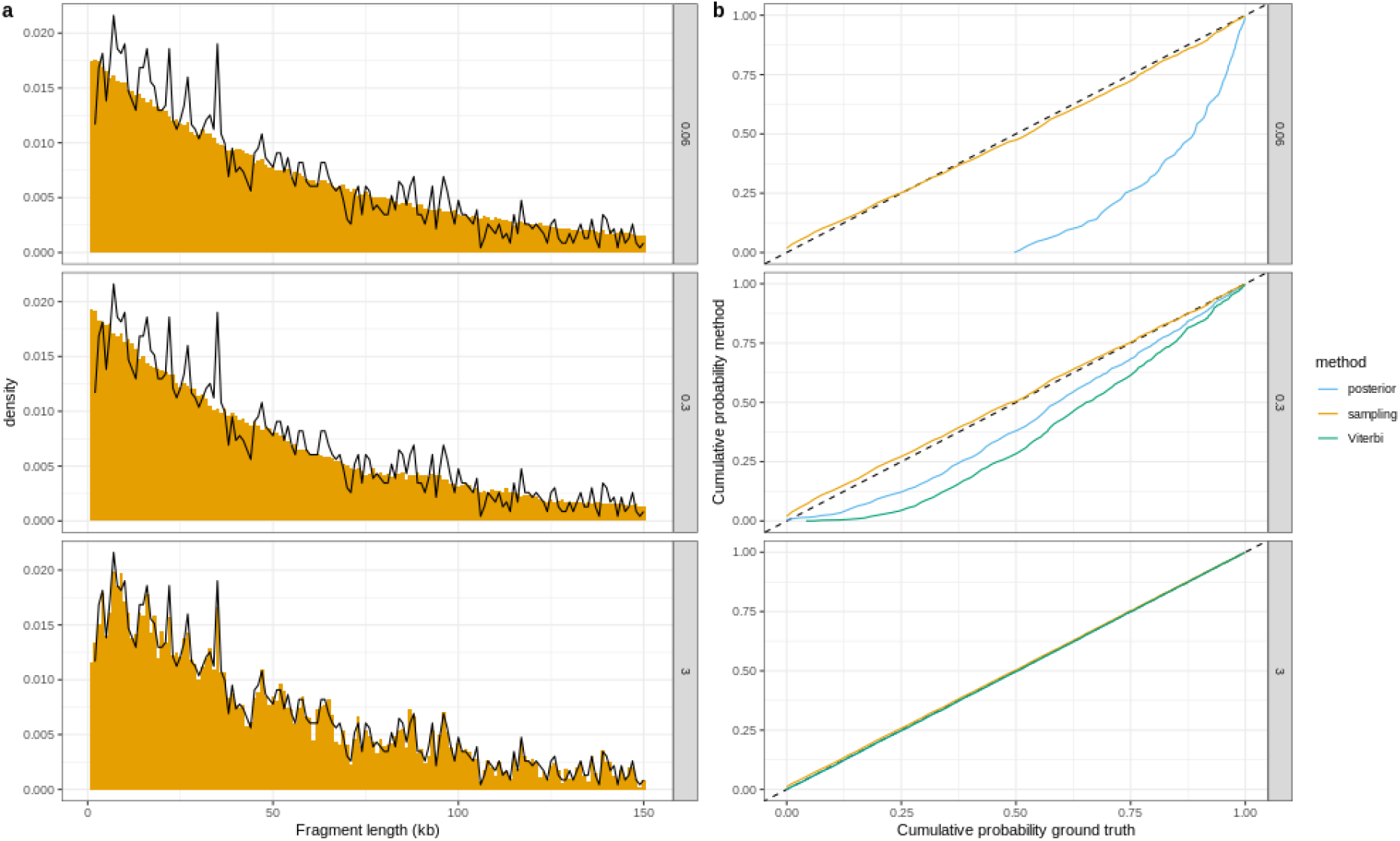
a) We show the true fragment length distribution in black for a 2.8 Gb genome with constant recombination rate. In orange we show the fragment length distribution from 100 inhomogeneous samples for three different *λ*_*A*_ values. b) we show the reverse cumulative probability density function (starting from the longest fragments). We only consider fragments shorter than 150 kb. For *λ*_*A*_ = 0. 06, Viterbi decoding never enters the archaic state.

We also explore varying recombination rate. We note that the true length distribution is skewed towards shorter fragments. For *λ*_*A*_ = 0. 3 which resembles the case of Neanderthal-human introgression, we are not able to recover the correct distribution with the sampling approach. This is due to many short fragments having 0 derived SNPs, which are impossible to detect.

However the length distribution obtained by the sampling approach is closer to the true distribution (Figure S6b). When *λ*_*A*_ = 3, all decoding methods can recover the correct length distribution.

**Figure S6.**
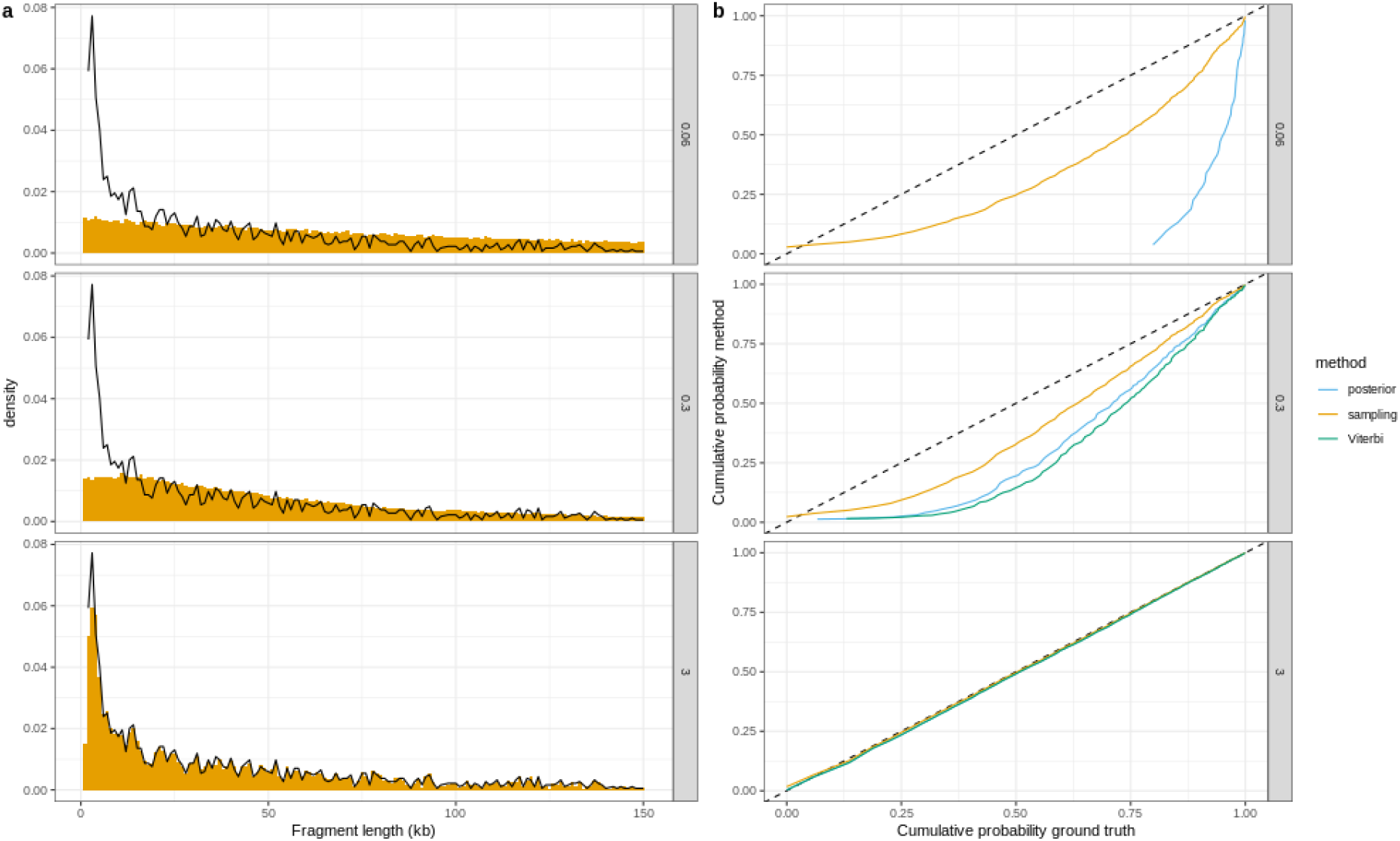
a) Comparison between the true (black line) and 100 samples from the posterior (orange histogram) fragment length distribution for a 2.8 Gb genome with varying recombination rate. b) The reverse cumulative probability density function (starting from the longest fragments). Only fragments shorter than 150 kb are considered. For *λ*_*A*_ = 0. 06, Viterbi decoding never decodes any archaic fragment.

### S3 *α* results are robust to variation for values within 95% CI

**Table S2.**
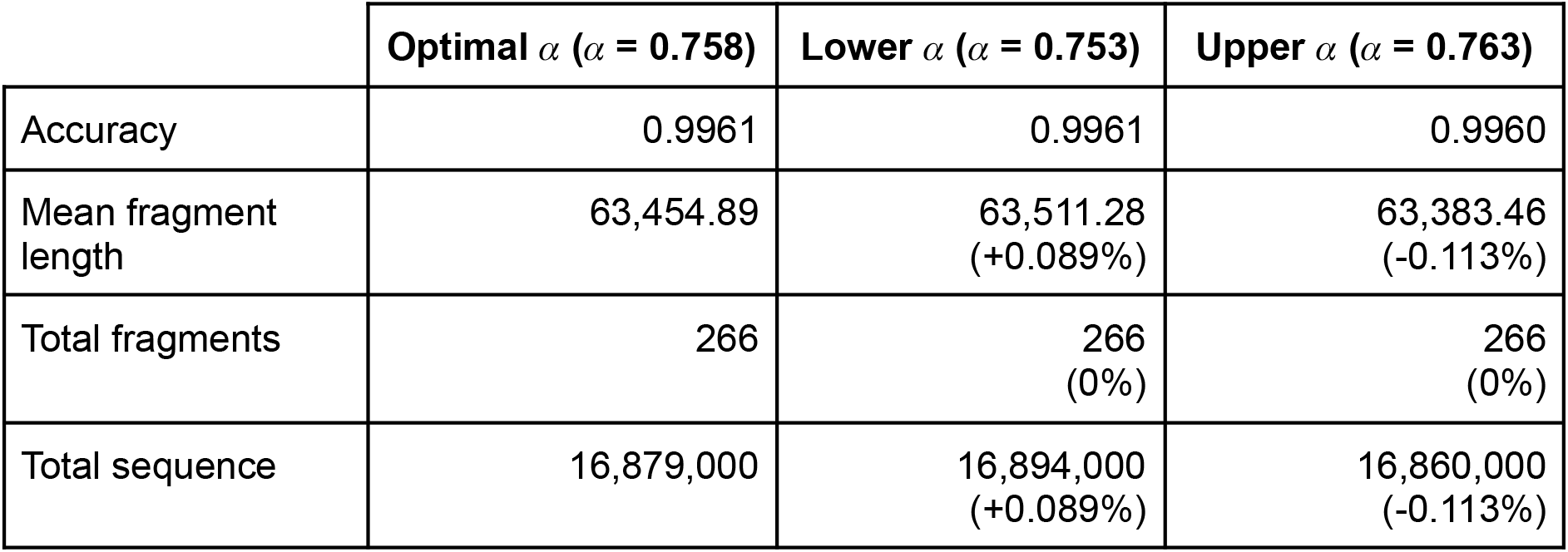
Comparison of relevant summary statistics for the hybrid paths using the optimal *α* and the lower and upper values of its 95% CI for the simulation based on realistic parameters. Difference to the optimal statistic is shown in percent.

|  | **Optimal $\alpha$ ($\alpha = 0.758$)** | **Lower $\alpha$ ($\alpha = 0.753$)** | **Upper $\alpha$ ($\alpha = 0.763$)** |
| --- | --- | --- | --- |
| Accuracy | 0.9961 | 0.9961 | 0.9960 |
| Mean fragment length | 63,454.89 | 63,511.28<br>(+0.089%) | 63,383.46<br>(-0.113%) |
| Total fragments | 266 | 266<br>(0%) | 266<br>(0%) |
| Total sequence | 16,879,000 | 16,894,000<br>(+0.089%) | 16,860,000<br>(-0.113%) |

### S4 Hybrid decodings compared to Viterbi and Posterior decoding on real data

**Table S3.** Number of archaic fragments that match (overlap for at least a base pair) between decoding methods per individuals’ haplotypes. Rows correspond to the method from which the target fragments have been obtained. Columns correspond to the method that the fragments are being compared to. For cells with the same method in the row and column, the value indicates the total number of archaic fragments inferred by that method. When the row is different than the column, the value corresponds to the number of fragments of the row method overlaping with the column method. If a cell is subdivided in three cells, the first entry indicates the number of fragments form the row method that overlap with a single fragment with the column method, the second the overlaps with two fragments and the third entry with three.

| NA19078 (hap1) | Viterbi | Hybrid |  |  | Posterior |  |  |
| --- | --- | --- | --- | --- | --- | --- | --- |
| Viterbi | 636 | 627 | 8 | 1 | 622 | 13 | 1 |
| Hybrid | 646 | 834 |  |  | 824 | 10 |  |
| Posterior | 651 | 844 |  |  | 964 |  |  |

| NA19078 (hap2) | Viterbi | Hybrid |  |  | Posterior |  |  |
| --- | --- | --- | --- | --- | --- | --- | --- |
| Viterbi | 633 | 622 | 11 | 0 | 614 | 18 | 1 |
| Hybrid | 644 | 836 |  |  | 825 | 10 | 1 |
| Posterior | 653 | 848 |  |  | 972 |  |  |

| NA20810 (hap1) | Viterbi | Hybrid |  |  | Posterior |  |  |
| --- | --- | --- | --- | --- | --- | --- | --- |
| Viterbi | 510 | 498 | 10 | 2 | 490 | 18 | 2 |
| Hybrid | 524 | 684 |  |  | 674 | 10 | 0 |
| Posterior | 532 | 694 |  |  | 783 |  |  |

| NA20810 (hap2) | Viterbi | Hybrid |  |  | Posterior |  |  |
| --- | --- | --- | --- | --- | --- | --- | --- |
| Viterbi | 542 | 529 | 13 | 0 | 520 | 20 | 2 |
| Hybrid | 555 | 729 |  |  | 719 | 9 | 1 |
| Posterior | 566 | 740 |  |  | 862 |  |  |

| NA21130 (hap1) | Viterbi | Hybrid |  |  | Posterior |  |  |
| --- | --- | --- | --- | --- | --- | --- | --- |
| Viterbi | 550 | 531 | 19 | 0 | 528 | 19 | 3 |
| Hybrid | 569 | 749 |  |  | 741 | 8 | 0 |
| Posterior | 575 | 757 |  |  | 884 |  |  |

| NA21130 (hap2) | Viterbi | Hybrid |  |  | Posterior |  |  |
| --- | --- | --- | --- | --- | --- | --- | --- |
| Viterbi | 535 | 519 | 16 | 0 | 515 | 19 | 1 |
| Hybrid | 551 | 729 |  |  | 719 | 10 | 0 |
| Posterior | 556 | 739 |  |  | 870 |  |  |

### S5 Linkage Disequilibrium

**Table S4.**
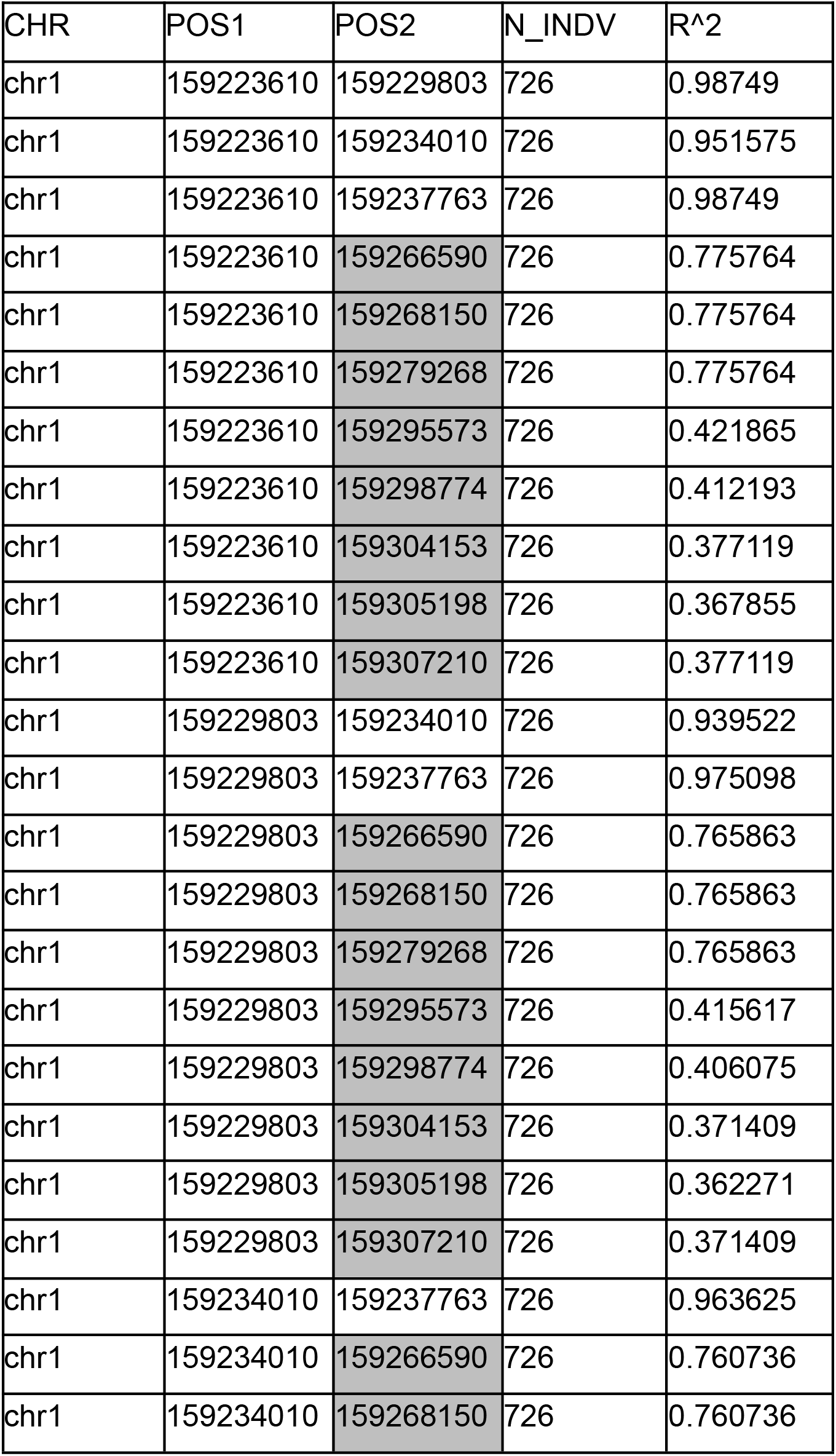
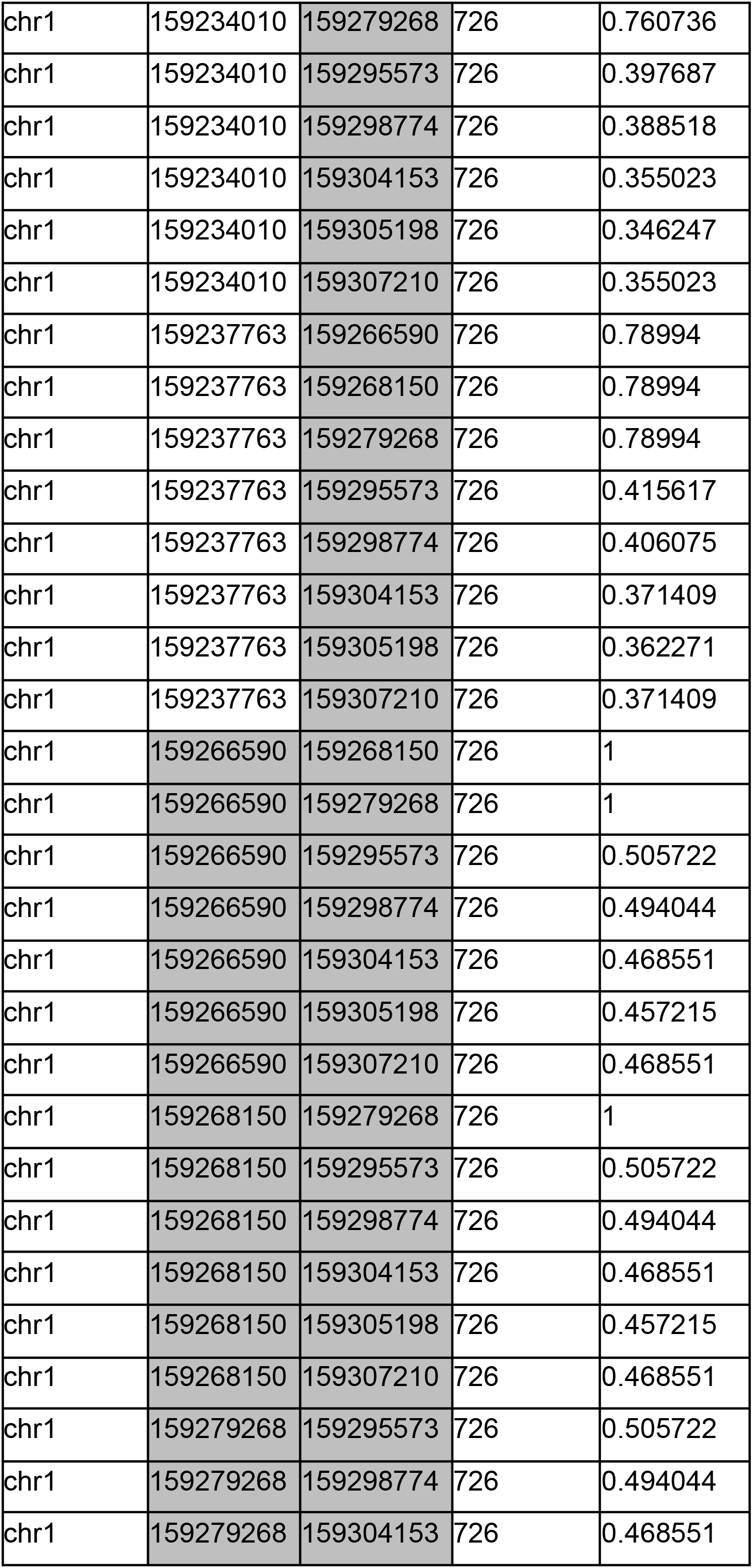
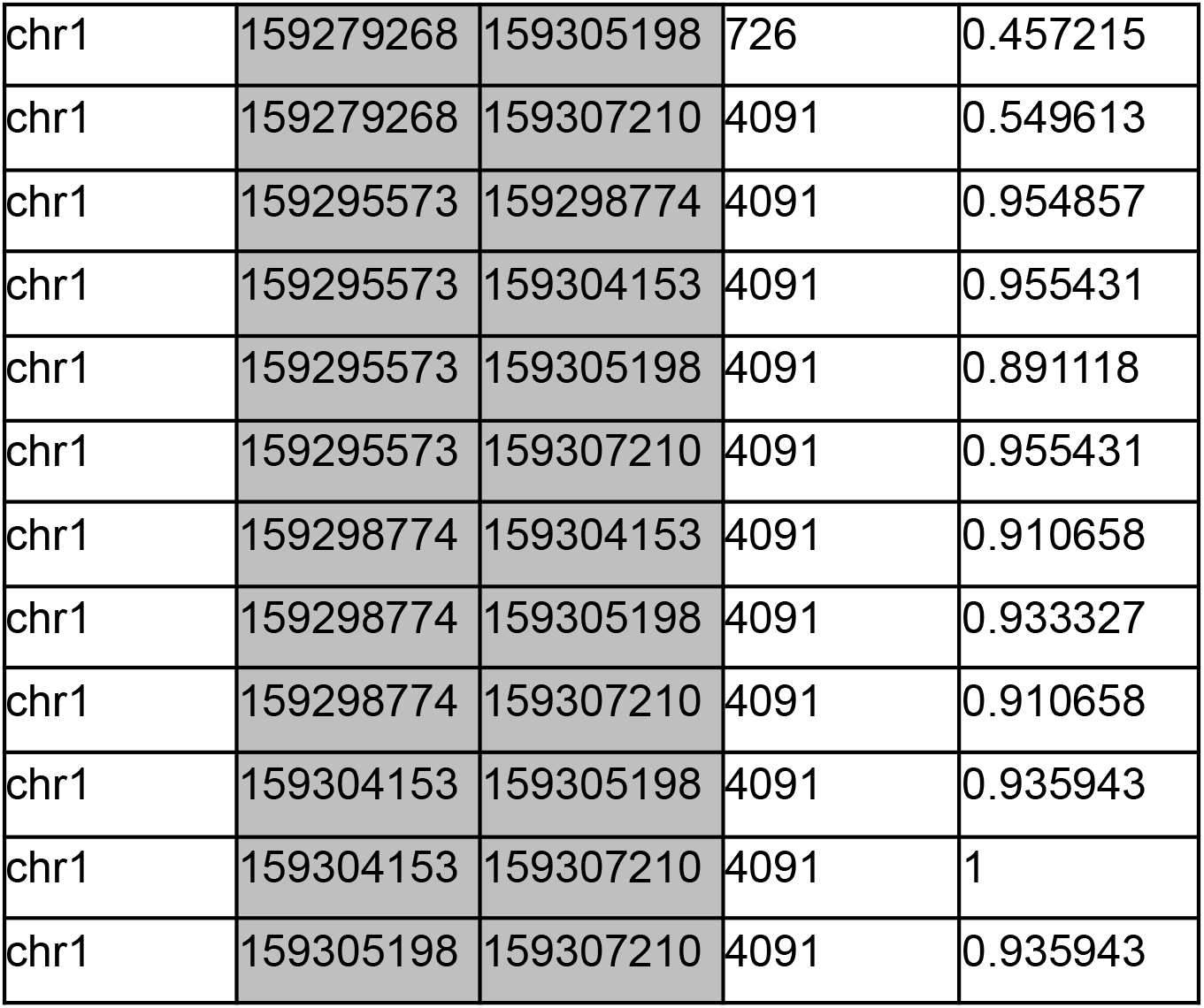
Linkage disequilibrim measured as *r*^2^ between all pairs of observations in the chr1:159203000-159329000 haplotype 1 of individual NA19078. All SNPs are also found in Neanderthal samples, except for SNP 159305198. SNPs identified in the second fragment called by Posterior decoding in Figure 7 are colored in grey.

## Bibliography

1. Aston, John A. D. and Donald E. K. Martin. 2007. Distributions associated with general runs and patterns in hidden Markov models. The Annals of Applied Statistics 1, Nr. 2: 585–611. doi:10.1214/07-aoas125,.

2. Bæk Zenia Elise Damgaard, Moisès Coll Macià, Laurits Skov and Asger Hobolth. 2025. Advanced posterior analyses of hidden Markov models: finite Markov chain imbedding and hybrid decoding. arXiv. doi:10.48550/arxiv.2504.15156,.

3. Baumdicker, Franz, Gertjan Bisschop, Daniel Goldstein, Graham Gower, Aaron P Ragsdale, Georgia Tsambos, Sha Zhu, et al. 2021. Efficient ancestry and mutation simulation with msprime 1.0. Genetics 220, Nr. 3: iyab229. doi:10.1093/genetics/iyab229,.

4. Bergström, Anders, Shane A McCarthy, Ruoyun Hui, Mohamed A Almarri, Qasim Ayub, Petr Danecek, Yuan Chen, et al. 2020. Insights into human genetic variation and population history from 929 diverse genomes. Science 367, Nr. 6484: eaay5012. doi:10.1126/science.aay5012,.

5. Coll Macià, Moisès, Laurits Skov, Benjamin Marco Peter and Mikkel Heide Schierup. 2021. Different historical generation intervals in human populations inferred from Neanderthal fragment lengths and mutation signatures. Nature Communications 12, Nr. 1: 5317. doi:10.1038/s41467-021-25524-4,.

6. Iasi, Leonardo N. M., Manjusha Chintalapati, Laurits Skov, Alba Bossoms Mesa, Mateja Hajdinjak, Benjamin M. Peter and Priya Moorjani. 2024. Neanderthal ancestry through time: Insights from genomes of ancient and present-day humans. Science 386, Nr. 6727: eadq3010. doi:10.1126/science.adq3010,.

7. Koenig, Zan, Mary T. Yohannes, Lethukuthula L. Nkambule, Xuefang Zhao, Julia K. Goodrich, Heesu Ally Kim, Michael W. Wilson, et al. 2024. A harmonized public resource of deeply sequenced diverse human genomes. Genome Research 34, Nr. 5: 796–809. doi:10.1101/gr.278378.123,.

8. Kuljus, Kristi and Jüri Lember. 2023. Pairwise Markov Models and Hybrid Segmentation Approach. Methodology and Computing in Applied Probability 25, Nr. 2: 67. doi:10.1007/s11009-023-10044-z,.

9. Lauterbur, M Elise, Maria Izabel A Cavassim, Ariella L Gladstein, Graham Gower, Nathaniel S Pope, Georgia Tsambos, Jeffrey Adrion, et al. 2023. Expanding the stdpopsim species catalog, and lessons learned for realistic genome simulations. eLife 12: RP84874. doi:10.7554/elife.84874,.

10. Lember, Jüri and Alexey A. Koloydenko. 2014. Bridging Viterbi and Posterior Decoding: A Generalized Risk Approach to Hidden Path Inference Based on Hidden Markov Models. Journal of Machine Learning Research. http://jmlr.org/papers/v15/lember14a.html.

11. Liang, Mason and Rasmus Nielsen. 2014. The Lengths of Admixture Tracts. Genetics 197, Nr. 3: 953–967. doi:10.1534/genetics.114.162362,.

12. Mallick, Swapan, Adam Micco, Matthew Mah, Harald Ringbauer, Iosif Lazaridis, Iñigo Olalde, Nick Patterson and David Reich. 2024. The Allen Ancient DNA Resource (AADR) a curated compendium of ancient human genomes. Scientific Data 11, Nr. 1: 182. doi:10.1038/s41597-024-03031-7,.

13. Perez, Gerardo, Galt P Barber, Anna Benet-Pages, Jonathan Casper, Hiram Clawson, Mark Diekhans, Clay Fischer, et al. 2024. The UCSC Genome Browser database: 2025 update. Nucleic Acids Research 53, Nr. D1: D1243–D1249. doi:10.1093/nar/gkae974,.

14. Petr, Martin, Svante Pääbo, Janet Kelso and Benjamin Vernot. 2019. Limits of long-term selection against Neandertal introgression. Proceedings of the National Academy of Sciences: 201814338. doi:10.1073/pnas.1814338116,.

15. Prüfer, Kay. 2018. snpAD: an ancient DNA genotype caller. Bioinformatics 34, Nr. 24: 4165–4171. doi:10.1093/bioinformatics/bty507,.

16. Sankararaman, Sriram, Swapan Mallick, Michael Dannemann, Kay Prüfer, Janet Kelso, Svante Pääbo, Nick Patterson and David Reich. 2014. The genomic landscape of Neanderthal ancestry in present-day humans. Nature 507, Nr. 7492: 354–357. doi:10.1038/nature12961,.

17. Sankararaman, Sriram, Swapan Mallick, Nick Patterson and David Reich. 2016. The Combined Landscape of Denisovan and Neanderthal Ancestry in Present-Day Humans. Current biology : CB 26, Nr. 9: 1241–7. doi:10.1016/j.cub.2016.03.037,.

18. Skov, Laurits, Ruoyun Hui, Vladimir Shchur, Asger Hobolth, Aylwyn Scally, Mikkel Heide Schierup and Richard Durbin. 2018. Detecting archaic introgression using an unadmixed outgroup. PLOS Genetics 14, Nr. 9: e1007641. doi:10.1371/journal.pgen.1007641,.

19. Skov, Laurits, Moisès Macià, Garðar Sveinbjörnsson, Fabrizio Mafessoni, Elise A Lucotte, Margret S Einarsdóttir, Hakon Jonsson, et al. 2020. The nature of Neanderthal introgression revealed by 27,566 Icelandic genomes. Nature: 1–6. doi:10.1038/s41586-020-2225-9,.

20. Vernot, Benjamin and Joshua M Akey. 2014. Resurrecting Surviving Neandertal Lineages from Modern Human Genomes. Science 343, Nr. 6174: 1017–1021. doi:10.1126/science.1245938,.

21. Zucchini, Walter, Iain L. MacDonald and Roland Langrock. 2017. Hidden Markov Models for Time Series, An Introduction Using R. doi:10.1201/b20790,.

